# Manipulating condensation of thermo-sensitive SUF4 protein tunes flowering time in Arabidopsis thaliana

**DOI:** 10.1101/2023.11.01.565081

**Authors:** Heather M. Meyer, Takashi Hotta, Andrey Malkovskiy, Yixian Zheng, David W. Ehrhardt

## Abstract

Intrinsically disordered proteins (IDP) lack stable tertiary structures, which allows them to change conformation and function under different physicochemical conditions. This may be highly advantageous for plants, which often use changes in their environment to elicit a variety of responses, including developmental events. For instance, some plants delay flowering in the fall and require exposure to winter temperatures as a cue to initiate flowering the following spring. Many of the genes involved in temperature-dependent flowering have been extensively studied in *Arabidopsis*, yet how plants coordinate their molecular states with seasonal temperature change is poorly understood. Here, we explore the role of temperature-sensitive phase separation of the IDP and flowering-time regulator, SUPPRESSOR OF FRIGIDA 4 (SUF4), in modulating flowering time. SUF4 has a well-defined role in regulating temperature-dependent flowering time by activating the master floral suppressor *FLOWERING LOCUS C (FLC)*. We show that in plant nuclei, SUF4 is a temperature-sensitive protein that assembles into biomolecular condensates in warm temperatures (20°C). When temperatures cool (4°C), SUF4 nuclear condensates disassemble, causing SUF4 to disperse within the nucleoplasm. Additionally, we demonstrate that the number of SUF4 condensates quantitatively correlates with flowering time. Progressive alterations to the amino acid composition of SUF4’s disordered region cause likewise progressive changes in temperature-dependent condensation both *in vitro* and *in vivo*, *FLC* transcription, and the onset of flowering. Lastly, we observe that SUF4 condensates coincide with the accumulation of other key flowering-time proteins (FRIGIDA and ELF7). These findings indicate that condensation of SUF4 likely plays a pivotal role in promoting flowering, possibly by concentrating and stabilizing regulatory factors needed for the transcriptional activation of *FLC* through temperature-dependent phase separation. This research suggests that in plants, IDPs can sense environmental cues and regulate critical developmental processes.

**One-Sentence Summary:** Evidence suggest that *Arabidopsis* uses temperature-sensitive condensation to regulate *FLC* transcription and flowering time.

## INTRODUCTION

The survival of all species depends on their ability to reproduce. For plants, reproductive success relies heavily on the onset of flowering, a phenological feature influenced by changes in seasons. Many plant species, including winter crops, have evolved to use long exposure to winter temperatures (e.g., vernalization) as a cue to flower in the following spring. Temperature-induced development plays a crucial role in the long-term reproductive success and yield of plants, enabling them to synchronize their flowering with seasonal pollinators and set seed during favorable climatic conditions. Thus, as climate change drives new spatial and seasonal temperature trends, it is increasingly important to understand how plants physically sense and integrate thermal shifts.

In the model plant *Arabidopsis thalian*a, temperature-dependent flowering is modulated by the epigenetic silencing of the *FLOWERING LOCUS C (FLC)*. *FLC* encodes a MADS-box transcription factor that represses flowering by silencing floral-inducing genes including *FT* and *SOC1*^1–3^. In warm temperatures, SUPPRESSOR OF FRIGIDA 4 (SUF4), FRIGIDA (FRI), and FRI-associated proteins transcriptionally activate *FLC* by recruiting chromatin modifiers, such as members of the PAF1 complex, which methylate histone H3 lysine 4 (H3K4me3) at *FLC*’s transcriptional start site ^3,4^. Conversely, exposure to cold temperatures promotes antisense long non-coding RNAs and a second set of chromatin remodelers belonging to the POLYCOMB REPRESSIVE COMPLEX 2 (PRC2) to epigenetically silence *FLC*, ultimately priming plants to flower the following spring ^5,6^. While many elements of the genetic and epigenetic regulation of temperature-dependent flowering are well established, the mechanisms by which plants sense and transduce thermal changes to elicit these responses remain a mystery.

Intrinsically disordered proteins (IDPs) are highly sensitive to changes in their physical environment ^7,8^. Unlike canonical proteins, which assume relatively stable structures with limited conformational changes in response to different stimuli and physiological conditions, IDPs possess low sequence complexity and a reduced ability to form stable tertiary structures. This inherent flexibility enables them to readily alter their conformation and function under varying physical and solution conditions. For example, many IDPs undergo phase separation into liquid-like protein droplets or viscoelastic hydrogels, facilitating rapid concentration, formation, and/or stabilization of higher-order complexes in response to environmental changes^8,9^. The ability for IDPs to phase separate has led to the intriguing hypothesis that this mechanism could serve as a biological means for cells and organisms to perceive and adapt to environmental signals^,10^. However, verifying this hypothesis poses a challenge due to the difficulty of distinguishing functions derived from phase separation from other protein properties.

Here, we identified that one of the key proteins required for regulation of *FLC* transcription and temperature-dependent timing of flowering, SUPPRESSOR OF FRIGIDA 4 (SUF4), exhibits significant similarities with BUGZ, an IDP known for its temperature-dependent phase condensation and involvement in mitotic chromosome alignment in animal cells^10–12^. Both proteins are comparable in size and feature two highly conserved zinc finger domains at their N-termini, followed by intrinsically disordered domains with low amino acid complexity enriched in proline, valine, and glycine residues. Given these structural parallels, along with BuGZ’s documented biophysical behavior and SUF4’s genetic and molecular implications in flowering timing, we asked if SUF4 also undergoes temperature-dependent phase transitions, using changes in its phase behavior to regulate the seasonal onset of flowering time.

Using confocal microscopy, genetics, and biochemistry, we discovered that SUF4 is indeed a temperature-sensitive protein that undergoes phase separation *in vitro*, forming proteinaceous droplets, and *in planta*, promoting the formation of nuclear condensates. At warm temperatures (20°C), SUF4 dynamically forms nuclear condensates, whereas at low temperatures (4°C), these condensates fail to form, causing SUF4 to disperse throughout the nucleoplasm. Furthermore, through use of targeted mutagenesis, we established the significance of SUF4’s disordered region in nuclear condensate formation and the regulation of flowering time. Plants expressing SUF4 variants with graded aromatic/hydrophobic-to-hydrophilic amino acid substitutions in this region exhibit graded alterations in condensate formation, *FLC* transcript levels, and flowering-time responses. To explore the relationship between SUF4 condensates and other proteins involved in flowering time, we labeled two significant *FLC* regulators, FRIDIGA (FRI), and the PAF1-complex member EARLY FLOWERING 7 (ELF7) with fluorescent markers. We observed that both proteins often appear in nuclear condensates that co-localize with a subset of SUF4 condensates, indicating that SUF4 condensation occurs where other important regulators of flowering time also accumulate. Interestingly, while FRI could assemble into condensates containing SUF4 in some tissues (lateral roots), its accumulation was apparently antagonized by SUF4 condensation in other tissue (primary roots). Our findings provide strong evidence supporting the hypothesis that SUF4 regulates flowering time by utilizing temperature-dependent phase separation, which likely functions to enhance the activity of regulatory factors, including itself, that are necessary for the transcriptional activation of *FLC*.

## RESULTS

### SUF4 forms nuclear condensates *in planta*

To investigate the hypothesis that SUF4 phase separation is important for the regulation of flowering time, we first needed to assess if SUF4 undergoes temperature-dependent condensation *in vivo*. To do this, we engineered a monomeric fluorescent mEGFP-SUF4 fusion protein expressed from the putative *SUF4* native promoter and 3’ UTR (pSUF4::mEGFP-SUF4) and introduced this construct into *suf4* (*FRI*) mutant plants. We identified several transgenic lines that rescued both the early flowering phenotype of the null mutant and the transcriptional activity of *SUF4* as measured by the transcripts level of its downstream target *FLC* (Supplemental Fig. 1A and 1B), suggesting that the *SUF4* transgene is fully functional and expressed at adequate levels. We picked one of these lines to perform all subsequent analyses regarding SUF4.

To assess the subcellular localization and behavior of SUF4 within living plant cells, we imaged live cells using confocal microscopy at room temperature. In both the shoot and the root, we observed SUF4 to exhibit a diffuse localization pattern within the nucleoplasm and accumulate in nuclear foci reminiscent of biomolecular condensates (Supplemental Fig. 2). Additionally, we found that SUF4 localized in the nucleolar cavity, an elusive membrane-less compartment that forms within the nucleolus (Supplemental Fig. 2). The nucleolar cavity is a thermo-responsive structure in plant cells that exchanges its contents with the nucleus ^13–15^. However, we directed our analysis toward the function of the nucleoplasmic foci outside of the nucleoli where most genes, such as *FLC*, are transcribed ^16,17^.

If SUF4 condensates function as thermosensors, their formation should exhibit temperature sensitivity across a physiologically relevant temperature range *in vivo*. To test this, we subjected SUF4 rescue plants to a 24-hour cold treatment at 4°C and then live imaged epidermal and cortical cells in the division and growth zones of primary roots as the plants rewarmed to 20°C. At the beginning of rewarming, mEGFP-SUF4 appeared diffuse and no condensates were detected (Fig. 1A and 1B). As plants began to warm up, SUF4 signal progressively partitioned into nuclear condensates, with condensates forming within seconds to minutes once exposed to room temperature conditions. To ensure that these changes were temperature-induced, we also imaged plants over the same time course under constant 20°C (Fig. 1B). Although some condensates showed evidence of forming and dissipating over the time scale of observation (<10mins), a portion of SUF4 always remained partitioned in condensates. Together, our results suggest that SUF4 is a temperature-sensitive protein that assembles into biomolecular condensates in warm temperatures and remains diffuse in the nucleoplasm in cold temperatures.

**Figure 1.**
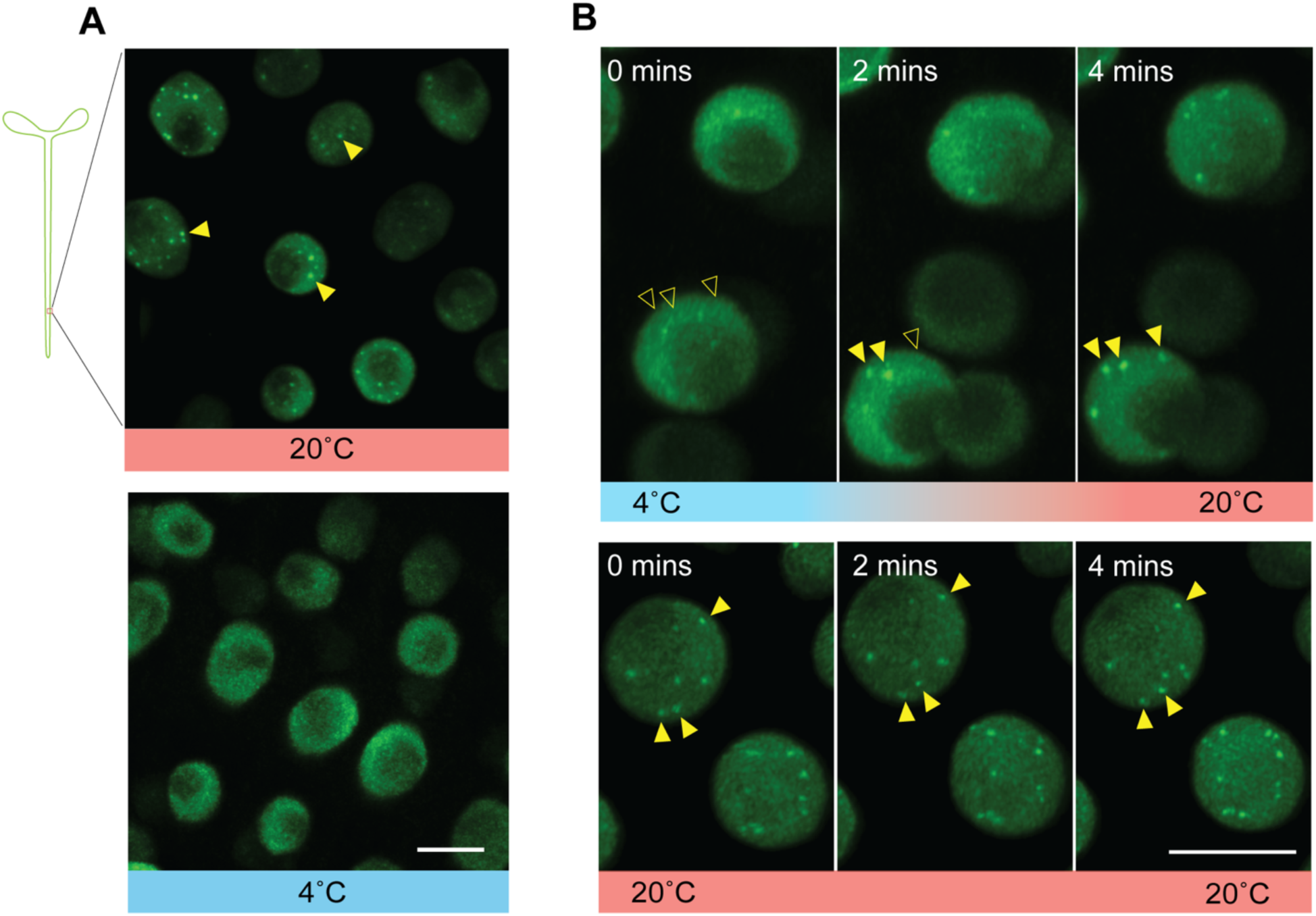
SUF4 forms nuclear condensates that are sensitive to temperature *in planta*. **A) mEGFP-SUF4 is expressed from native expression sequences in a null mutant background, *suf4* (FRI).** mEGFP-SUF4 is diffuse within the nucleoplasm and forms nuclear condensates at room temperature. However, at 4°C, condensates do not form, leaving SUF4 diffused in the nucleoplasm. **B)** mEGFP-SUF4 rapidly forms condensates as temperatures increase. Scale bar: 10µm, yellow arrowheads point to the locations of nuclear condensates (filled) or locations where they will appear in later images (empty).

### SUF4 phase separates *in vitro*

SUF4 is orthologous to the animal protein BuGZ (Supplemental Fig. 3), an IDP initially found to undergo temperature-dependent phase separation *in vitro* and regulate spindle assembly *in vivo*^10–12^. Therefore, to assess the potential for SUF4 to undergo phase separation independently of any plant cell factors, we expressed and purified 6xHis-mVenus-SUF4 (isoform 1, WT) from SF9 insect cells. Using both fluorescence and DIC microscopy, we observed that 6xHis-mVenus-SUF4 underwent phase separation at high temperatures (25<u>µM</u> at 37°C) and remained miscible at low temperatures (25<u>µM</u> at 4°C) when suspended in physiologically relevant buffers (Fig. 2A, Supplemental Fig. 4, see methods). These results suggest that, similar to BuGZ, SUF4 is a phase-separating protein displaying low critical solution temperature (LCST) behavior. To investigate further the phase behavior of SUF4, we conducted a turbidity assay to determine the thermal setpoint required for SUF4’s condensation. We found that 25µM of 6xHis-mVenus-SUF4 was fully miscible between 4-10°C but became increasingly turbid as temperatures gradually rose to 30°C (Fig. 2B). Additionally, in line with other phase-separating proteins, SUF4’s phase transition is concentration-dependent, with lower concentrations of SUF4 requiring higher temperature regimes to undergo phase separation (Fig. 2B, Supplemental Fig. 4). These results indicate that, similar to BuGZ, SUF4 is a temperature-sensitive protein that undergoes LCST phase separation *in vitro*.

**Figure 2.**
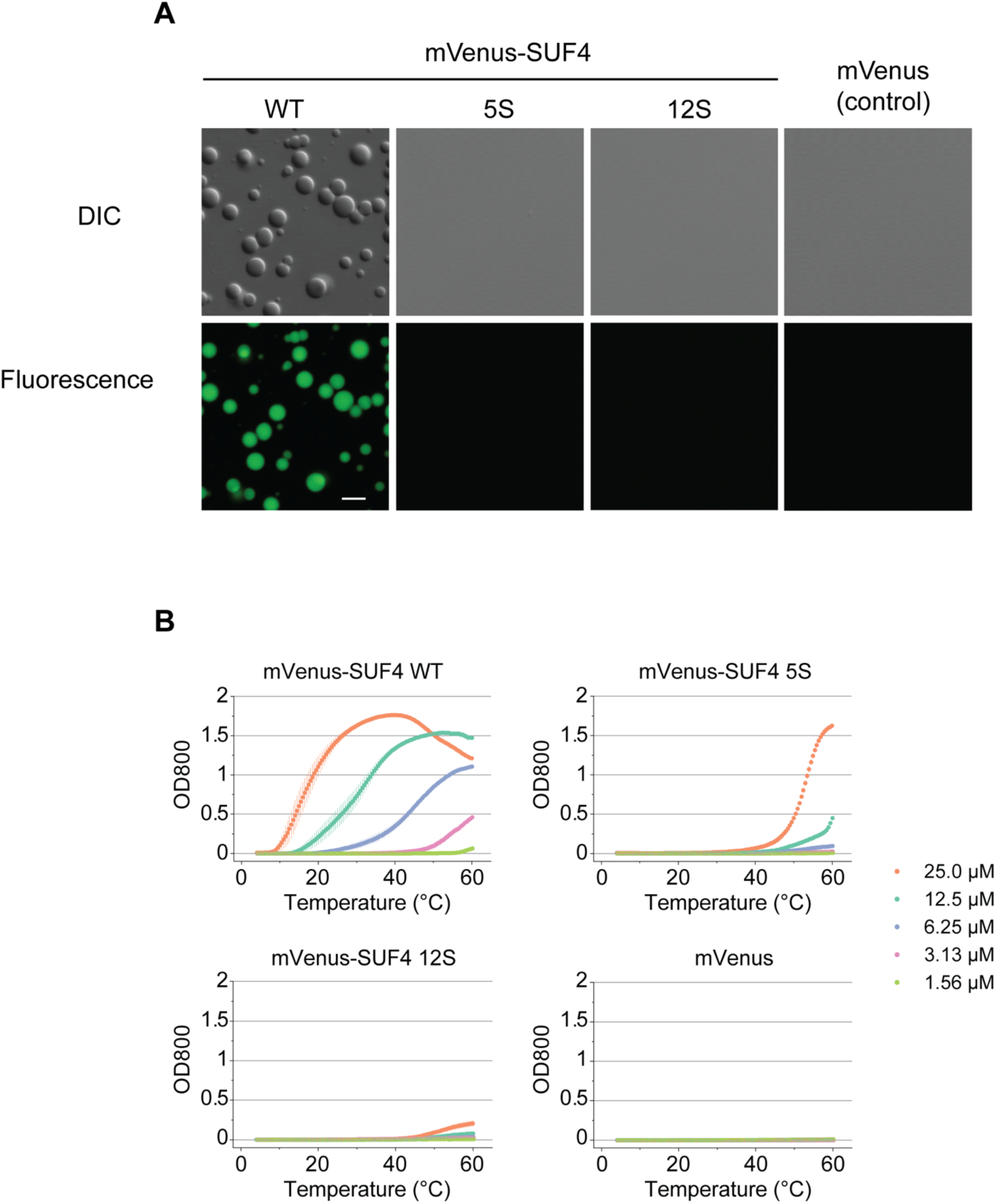
SUF4 forms temperature-sensitive condensates that may be tuned *in vitro*. **A)** DIC and confocal images of purified SUF4 variants. Only rescue SUF4 forms condensates at 37°C. Scale bar:5µm. B) Turbidity assay demonstrating the temperature sensitivity of purified SUF4 variants. The thermal set points shift depending on the severity of amino acid substitutions and the concentration used. Each graph includes three replicates for each sample, error bars depict standard errors.

### SUF4’s disordered region drives nuclear condensate formation *in vitro* and *in planta*

IDP phase separation is driven by multivalent interactions, such as those facilitated by aromatic/hydrophobic residues like tyrosine and phenylalanine. Moreover, these same residues are required for *in vitro* condensation of SUF4’s ortholog, BuGZ^11,12^. Despite being a proline-rich protein, SUF4 contains a total of 14 regularly spaced tyrosine and phenylalanine residues within its disordered region, with two of these residues falling in putative splice sites (Supplemental Fig. 3). To test if aromatic/hydrophobic residues in SUF4’s disorder region promote its temperature-dependent condensation, we created a SUF4 variant, termed 12S SUF4, in which we substituted 12 non-splice-interfering tyrosine and phenylalanine residues with serine and asked if this variant had a different phase separation thermal setpoint *in vitro*. We found 12S SUF4 had a large shift in its thermosensitive capabilities to phase separate, such that 25µM of purified 12S SUF4 now required temperatures nearing 50°C to begin phase separating as opposed to 20°C for the native protein (Fig. 2B). These results suggest that changing SUF4’s aromatic/hydrophobic patterning is sufficient to disrupt condensate formation *in vitro* by increasing the thermal set point at which SUF4 can phase separate.

To test if changes in SUF4 aromatic/hydrophobic patterning also influenced condensate formation *in vivo*, we generated the same 12 mutations in the mEGFP-SUF4 fusion protein expressed over a chromosomal null locus (pSUF4::mEGFP-12S SUF4). Similar to our observations *in vitro,* we found that 12S SUF4 exhibited a severe reduction in condensate formation *in vivo*. On average, 12S SUF4 plants formed 0-1 nuclear condensates per nucleus in root epidermal and cortical nuclei at 20°C, as compared to 4-5 per nucleus formed in rescue SUF4 (Fig. 3A and 3B, nuclei observed: 12S SUF4 n= 27, rescue SUF4 n=32). These results suggest that the altered coacervation properties of 12S SUF4 are maintained in the complex environment of the living cell.

**Figure 3.**
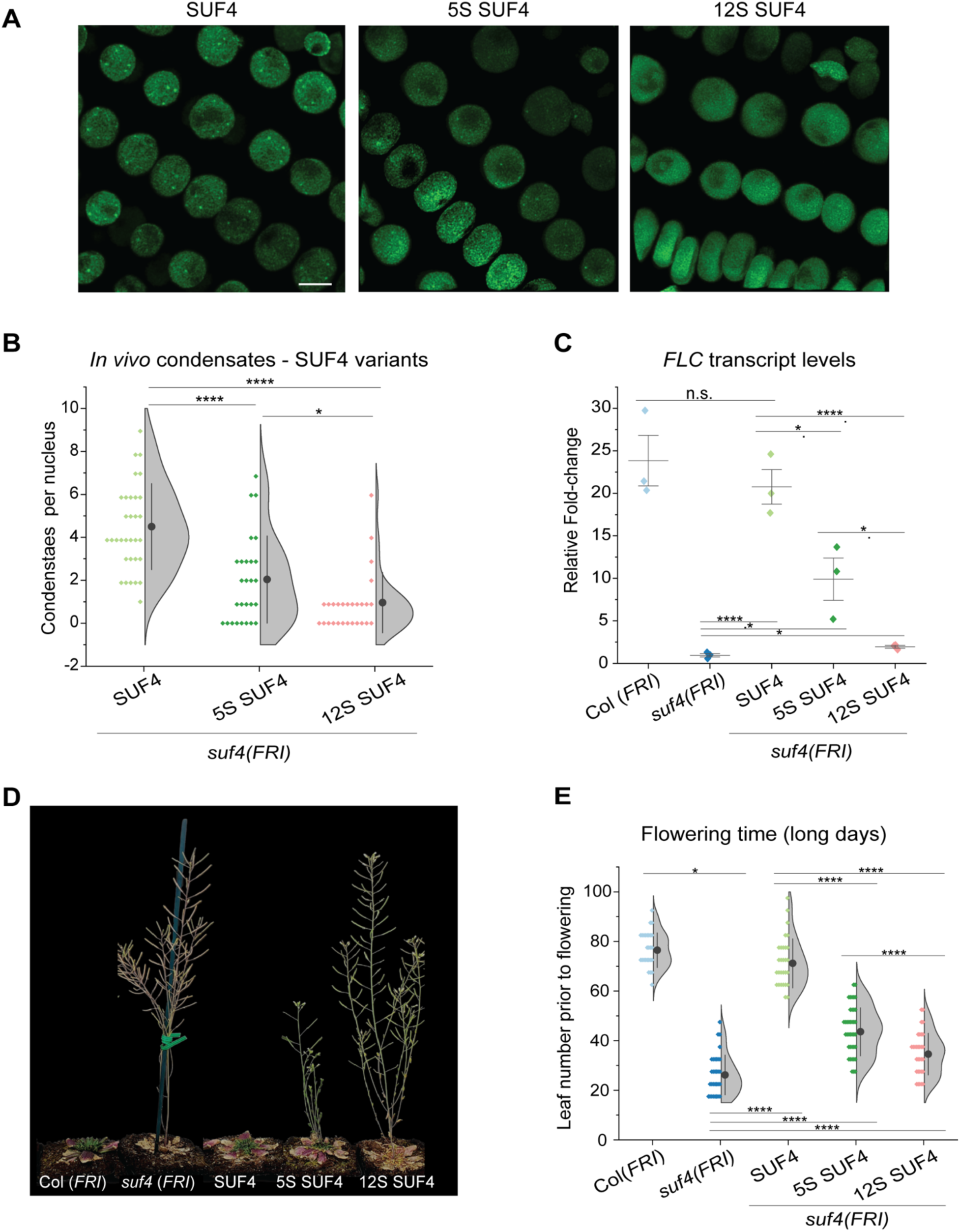
Tuning SUF4 condensate behavior corresponds to changes in flowering time outcomes. **A)** Confocal images of SUF4 variants, illustrating varying condensation patterns. Rescue mEGFP-SUF4 forms an average of 4-5 nuclear condensates; 5S SUF4 forms 2-3 nuclear condensates on average; 12S SUF4 shows 0-1 condensates on average. Scale bar: 5µm. **B)** Quantification of panel A. Statistical analysis was performed by Mann-Whitney U test. **C)** Relative abundance of *FLC* transcripts in different SUF4 variants. Rescue SUF4 shows *FLC* transcript levels similar to WT, while various substitutions lead to graded reductions. Statistical analysis conducted using a two-tailed Student’s t-test for independent means. Error bars represent the standard deviation. **D)** Image comparing the flowering time phenotypes of plants expressing various mEGFP-SUF4 variants with those of wildtype Col *(FRI)* plants. From left to right: Col *(FRI)* showing delayed flowering, *suf4* (FRI) mutant displaying early flowering, rescue mEGFP-SUF4 (functional copy of SUF4) expressed in *suf4 (FRI)* restoring the delayed flowering phenotype, 5S SUF4 (5 Y and F substitutions for S) expressed in *suf4 (FRI)* exhibiting a weak early flowering phenotype, and 12S SUF4 (12 F and Y substitutions for S) expressed in *suf4 (FRI)* showing a moderate early flowering phenotype. **E)** Quantification of panel D. Statistical analysis performed by Mann-Whitney U test. Asterisks denote specific p-values * <0.01, ** <0.001, ***<0.001, and ****<0.0001. Error bars are standard deviation.

Next, we investigated if SUF4 uses phase separation as a means to regulate flowering time. As a first step in testing this hypothesis, we asked if 12S SUF4 variant plants displayed a phenotypic difference in its response as compared to the rescue SUF4 plants. Using a leaf counting assay, we found that 12S SUF4 plants transitioned to flowering much sooner, producing significantly fewer rosette leaves before flowering (Long Day (LD): 35 leaves (n=30)) as compared to SUF4 rescue plants (LD: 71 leaves (n=29)), but significantly more so than the *suf4* null mutant (LD: 26 leaves (n=31)) (Fig. 2C-E, Supplemental Fig. 5). Previous research has shown that high levels of *FLC* must be produced for the suppression of flowering time. We therefore asked if the flowering time phenotype caused by the 12S mutant protein was correlated with a reduction in *FLC* transcripts. We found plants expressing the 12S variant to produce approximately 18-fold fewer *FLC* transcripts than our SUF4 rescue lines but 2-fold more than *suf4* null mutants (Fig. 3C). Thus, although *FLC* still undergoes some transcription in the 12S SUF4 variant, the mutations that caused disruption of SUF4 temperature-sensitive condensation greatly accelerated the onset of flowering.

### Tuning SUF4 phase behavior causes graded changes in *FLC* transcription and flowering time

Previous work from Jiang et al (2015) has shown that the *in vitro* phase behavior of SUF4’s ortholog, BuGZ, may be tuned by incrementally decreasing the number of combined phenylalanine and tyrosine to serine substitutions^10^. This finding, paired with the correlation between SUF4 phase separation and flowering time, prompted us to further assess if SUF4’s phase change properties themselves were critical for dictating different flowering time outcomes. We reasoned that if so, substituting a smaller number of the phenylalanine and tyrosine residues should lead to a scalable intermediate phase separation phenotype and proportionally tune flowering time outcomes via decreasing the transcriptional levels of *FLC*. To test this prediction, we created an intermediate SUF4 variant that only had 5 combined tyrosine and phenylalanine to serine substitutions termed, 5S SUF4, and assessed its phase separating thermal setpoint *in vitro*. Consistent with what had been observed for BuGZ, we found purified 5S SUF4 to have a thermal shift that ranges between the phase change of rescue SUF4 and 12S SUF4 *in vitro* (Fig. 2B; 25µM showed a steep phase transition at 40° C instead of 20°C like rescue SUF4 and 50°C like 12S SUF4).

To test the function of the 5S SUF4 variant *in vivo*, we introduced it into *suf4* null mutant plants and asked if these plants exhibited intermediate phenotypes relating to SUF4 phase separation, *FLC* transcriptional output, and flowering time. Remarkably, we found 5S SUF4 plants to display an intermediate condensate phenotype *in vivo*, in which they formed significantly fewer nuclear condensates in root epidermal and cortical cells than rescue SUF4 but more than 12S SUF4 (Fig. 3A and 3B; rescue SUF4:4-5, 5S SUF4:2-3, 12S SUF4:0-1). This outcome differed from the results observed with 5S SUF4 purified protein, which showed no phase separation at room temperature, suggesting that the complex cellular environment may influence the phase behavior of 5S SUF4 (Figure 2A). Moreover, 5S SUF4 plants exhibited *FLC* transcript levels and flowering time responses that both fell between rescue SUF4 and 12S (Fig. 3C-E, Supplemental Fig. 5; rescue SUF4 produced 20-fold/ LD: 71 leaves (n=29), 5S SUF4: 10-fold/LD: 43 leaves (n=39), 12S SUF4 3-fold/ LD: 35 leaves (n=30)). These results indicate that tuning the LCST behavior of SUF4 is sufficient to predictably change the onset of flowering through modulating transcript levels of *FLC*.

### SUF4’s exchange dynamics scale with changes in LCST behavior

Biomolecular condensates may assume a broad range of material properties, spanning liquid-, gel-, or solid-like structures, with these respective properties often being hypothesized as being important for their biological function^7,18–20^. Under *in vitro* conditions, purified and condensed SUF4 exhibited behavior consistent with a liquid, similar to its ortholog BuGZ. For example, *in vitro* condensates on glass assume a round outline, as expected at a liquid-liquid interface, and are rapidly and repeatedly reversible with temperature change (Fig. 2A, Supplemental Fig. 4). However, whether SUF4’s material properties remain liquid-like during interactions with other biomolecules under native cellular conditions remains unknown. Previous research has suggested that the exchange dynamics of molecules within condensates are expected to correlate with their material state, with faster exchange rates in liquid-like condensates and slower rates in more solid-like structures^20,21^. Thus, to investigate the exchange dynamics of SUF4 and whether these dynamics are affected in our SUF4 variants, we conducted fluorescence recovery after photobleaching (FRAP) experiments on rescue SUF4, 5S SUF4, and a hydrophobic variant termed 30I SUF4, which contains 30 alanine-to-isoleucine substitutions in its disordered region (Supplemental Fig. 6). We excluded the 12S SUF4 variant as it rarely formed condensates. FRAP was performed in a small region of approximately 1.094 microns^2^, centered on a prominent mEGFP-SUF4 nuclear body in each cell assayed. Our findings revealed that changing the hydrophobicity of SUF4 greatly affected its FRAP dynamics. 30I SUF4 condensates did not reach their half recovery time within a 10-minute period, whereas both rescue SUF4 and 5S SUF4 rapidly reached their half recovery time within seconds (Fig. 4A, rescue SUF4: t1/2=1.27 seconds, n=13; 5S SUF4 t1/2=0.85 seconds, n=12, 1-sided t-test p<0.01). Additionally, 30I SUF4 condensates exhibited a higher “immobile” fraction compared to SUF4 rescue and 5S SUF4, which both largely recovered (30I SUF4 = 0.094 mobile fraction, SUF4 rescue = 0.85 mobile fraction, and 5S SUF4 = 0.76 mobile fraction; Fig 4A). Interestingly, despite 30I SUF4 having formed many condensates that were relatively slow to recover, we did not observe any rescue in plant phenotype concerning *FLC* transcription or flowering time (Supplemental Fig. 6). Previous studies have suggested that the biophysical properties of some condensates must fall within a fluidic window, in which being overly solid or fluid-like can lead to a decrease in functionality ^20–25^ . Therefore, in agreement with these findings, we propose that SUF4 needs to maintain a specific state of fluidity in order to function optimally.

**Figure 4.**
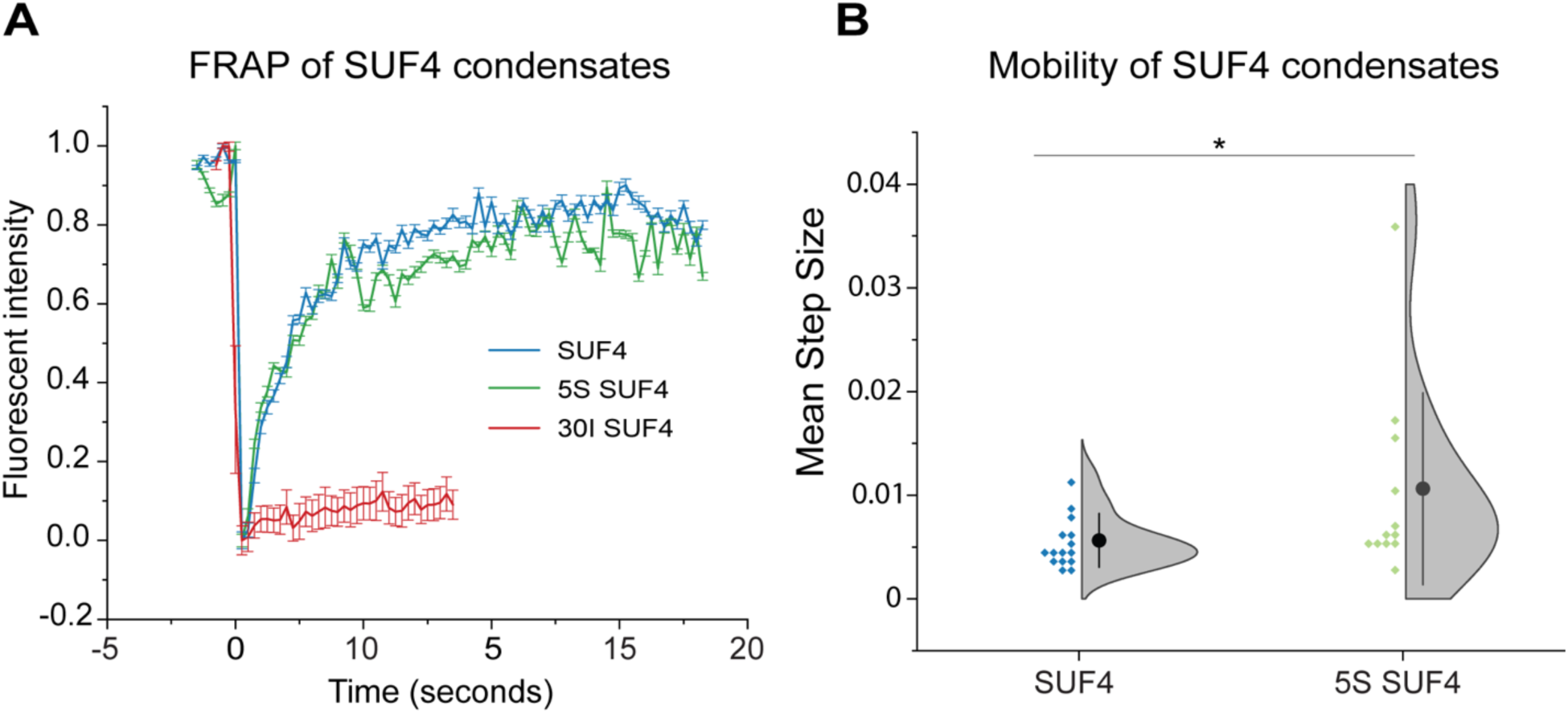
SUF4 variants exhibit altered exchange rate and mobility dynamics. **A)** Fluorescence Recovery After Photobleaching (FRAP) analysis of SUF4 variants reveals different recovery rates. 5S SUF4 exhibits the fastest recovery rate (t1/2=0.85 seconds), followed by rescue SUF4 (t1/2=1.27 seconds). 30I SUF4 does not recover within the duration of the experiment. **B)** Quantification of the total distance traveled for condensates used in panel A). On average 5S condensates were more mobile. Asterisks denote P-values from Student’s T-test * <0.01 FRAP distributions between rescue and 5S SUF4 were assessed for significance using a two-tailed Student’s t-test (p-value=0.007). Error bars= standard deviation.

A notable finding from our FRAP experiments is that the recovery curves of 5S SUF4 signal exhibited a lower R^2^ value on average when fitted to a single exponential recovery model, along with a lower bleaching depth compared to those observed in our rescue lines (rescue SUF4 R^2^ = .73, 5S SUF4 R^2^ = .53). This observation prompted us to further investigate the behavior of condensates in this background. In doing so, we noticed that 5S SUF4 condensates appeared to display greater motility compared to rescue SUF4 condensates. An increase in particle motility could explain the reduced model fit as movement of condensates out of the focal plane would tend to reduce their brightness, thus affecting measurement of signal recovery kinetics. To quantify the perceived difference in motility, we measured the total distance traveled by each condensate during our FRAP analysis. On average, we found that 5S SUF4 condensates traveled significantly greater distances within the nucleoplasm compared to rescue SUF4 condensates (Fig. 4B, average distance of 1.802 microns for rescue SUF4 and 3.046 microns for 5S SUF4). These results indicate that 5S SUF4 condensates are less constrained in their positional behavior compared to their rescue SUF4 counterparts.

### SUF4 colocalizes with FRI within nuclear condensates at room temperature in root cells recently derived from meristems

SUF4 has been shown previously to physically and genetically interact with FRIGIDA (FRI) and other members of the vernalization pathway, and this complex has been proposed to transcriptionally activate *FLC* ^3,4^. Therefore, to ask if SUF4 condensation occurs where other flowering time regulators are present, and to explore the possibility that SUF4 condensation might function to regulate *FLC* by concentrating flowering time proteins, we tested if there was an observable spatial relationship between SUF4 and FRI *in vivo*. We generated a fluorescently-labeled FRI fusion gene (FRI-mScarletI) that expresses from the native upstream sequence of FRI (pFRI) and transformed it into a mEGFP-SUF4 rescue line. Under room temperature conditions, we found FRI-mScarletI to exist in one of two distinct phase states within the nucleoplasm of root epidermal and cortical cells within the division and growth zones— either uncondensed and diffuse within the nucleoplasm or assembled into one-to-two apparent condensates (Fig 5A and B).

**Figure 5.**
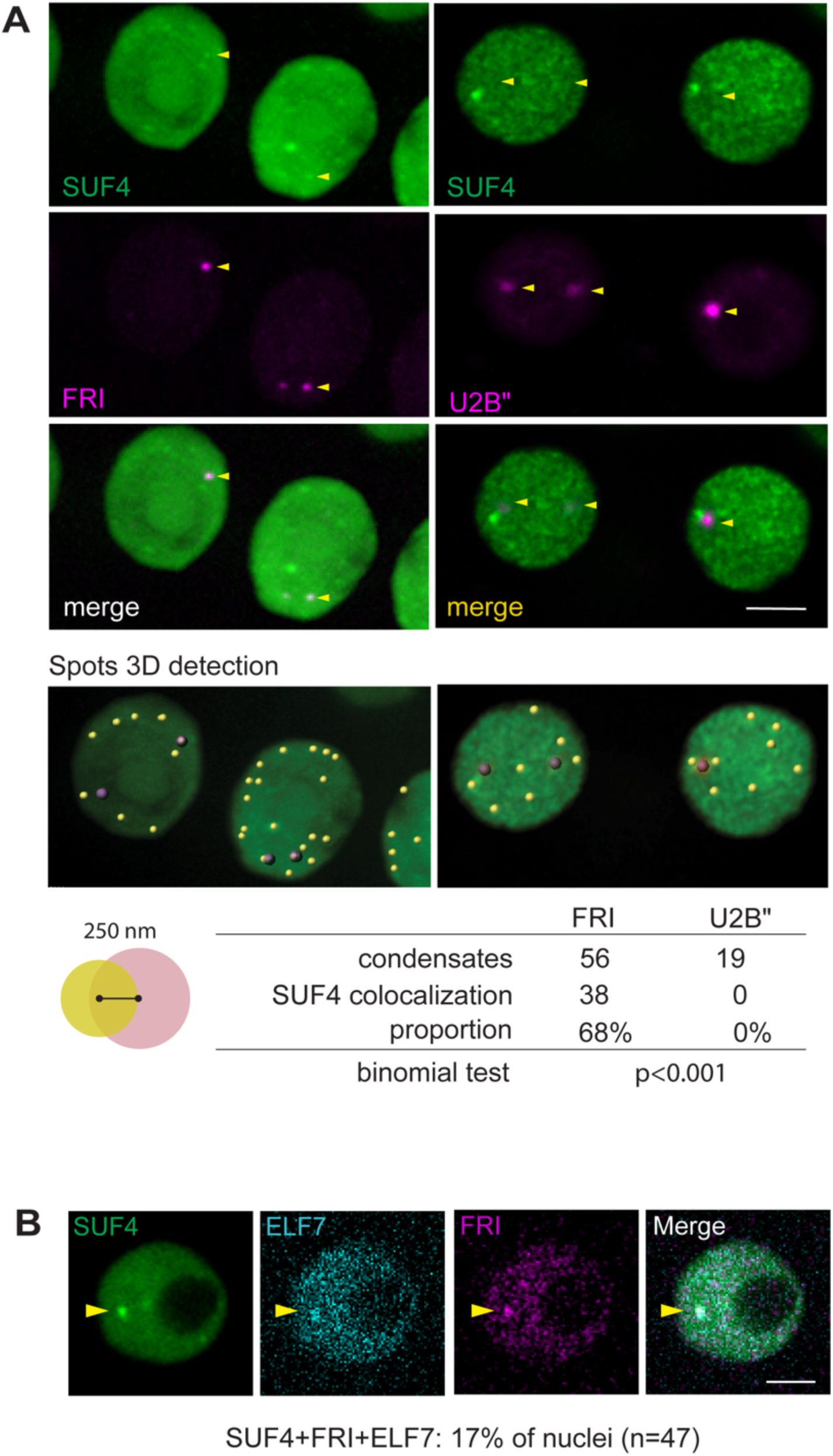
In warm temperature conditions, SUF4 co-localizes with members of the vernalization complex in newly derived meristematic tissues. **A)** SUF4 and FRI co-localize in nuclear condensates within developing lateral roots at room temperatures. FRI is visualized by expression of mScarletI-FRI. mScarletI-FRI co-localization appears to be selective, as U2B”, a protein found in nuclear splicing speckles, does not co-localize with SUF4. U2B” is visualized by expression of U2B”-mRFP. Binomial tests of proportions used to test for statistical significance. Arrowheads point to locations of tagged FRI or U2B” puncta. **B)** ELF7, a chromatin remodeler that functions with SUF4 and FRI to transcriptionally activate *FLC,* is visualized as mTFP-ELF7. mTFP-ELF7 also co-localizes within SUF4-FRI condensates. Arrowheads point to a nuclear body with signal concentration in all three channels. Scale bars: 5µm

Next, we tested if FRI-mScarletI condensates colocalized with mEGFP-SUF4 condensates. To measure co-localization, we considered condensates to be colocalized if the centroids of spot-detected condensates were closer together than the radius of the kernel used to detect the SUF4 condensates (<250nm). Out of 56 FRI-mScarletI condensates analyzed, 38 of them co-localized with a SUF4 condensate (Fig. 5A; 68% co-localization, n=47 nuclei, distance threshold=0.3µm). To ask whether this degree of colocalization was specific to the vernalization complex and not likely due to chance alone given the density of SUF4 nuclear condensates, we co-transformed the nuclear splicing speckle marker U2B”-mRFP into mEGFP-SUF4 rescue plants, a marker, like that for FRI, produces 1-2 bodies of larger size per nucleus. In the 18 nuclei examined, we found no instances of U2B’’ bodies co-localizing with SUF4 at room temperature conditions (0/19 U2B” speckles; Fig. 5A). These data demonstrate a remarkable degree of colocalization between SUF4 and FRI, suggest that the formation of FRI condensates under warm temperatures is not a random occurrence but rather occurs preferentially where condensation of SUF4 also occurs.

### SUF4 and FRI colocalization depends on tissue type and temperature

FRI has recently been reported to have no quantifiable physical interactions with SUF4 and to accumulate into nuclear condensates when exposed to extended periods of low temperatures ^26–28^. Thus, to test the spatial relationship between SUF4 and FRI in the cold, we exposed mEGFP-SUF4 and FRI-mScarletI co-expressing plants to 4°C for 24 hours and imaged their root cells as the plants warmed up. Consistent with previous findings^26–28^, we found FRI to form cold-induced nuclear condensates. However, unlike the case at room temperature (∼21°C), these condensates did not co-localize with SUF4. Instead, SUF4 condensates would sometimes appear in the same locations where FRI condensates previously existed as plants began to warm (observed in at least 7 nuclei; Fig 6A and B). These results suggest an antagonistic or independent relationship between cold-induced FRI condensates and SUF4. Additionally, they suggest that FRI undergoes cold-induced phase separation that is independent of *FLC* activation (a process reliant on SUF4).

**Figure 6.**
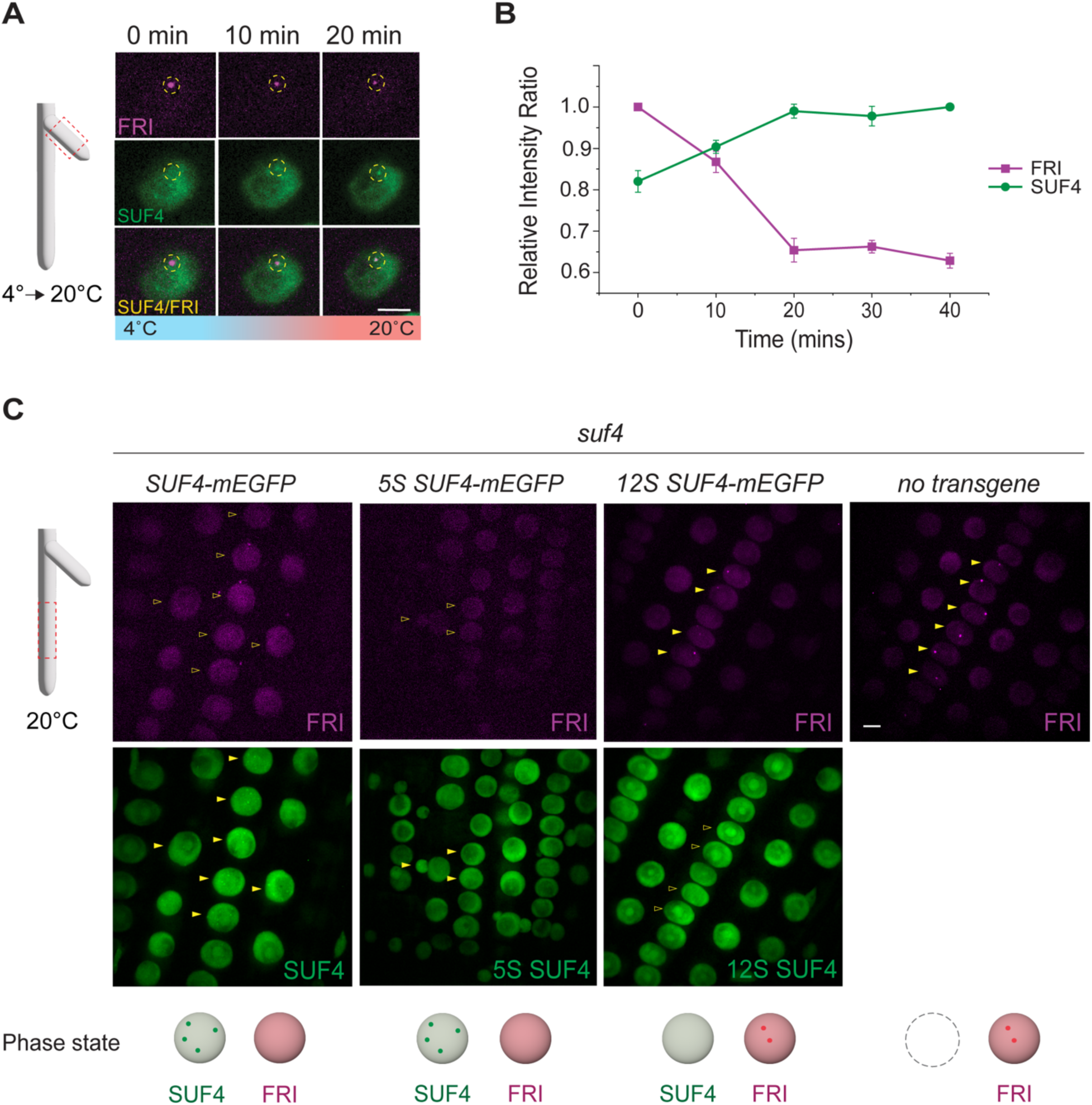
SUF4 and FRI have different localization patterns under cold temperatures. **A and B)** Cold-induced FRI condensates do not co-localize with SUF4. However, these condensates can exchange positions with SUF4 nuclear condensates upon warming. **A)** The first 20 minutes of an imaging series, showcasing the phase change behavior of FRI and SUF4 as temperatures rise. Dotted circle is not the ROI used for quantification but used to mark diminishing and appearing condensates. **B)** Graph quantifying the opposing behavior of FRI and SUF4 condensation state at four locations from the same times series shown in A) (40 minutes, five time points). Error bars are standard errors. **C)** SUF4 condensates spatially limit the formation of FRI condensates in the primary root. In SUF4 variant lines lacking condensates (12S and *suf4* mutants), FRI-mScarletI condensates begin to emerge. Schematic representing the phase state of SUF4 (green circles) and FRI (magenta circles) in nuclei of each genotype. Scale bars in all panels: 5µm.

Given the established relationship between the spatiotemporal organization of proteins and their biological function, we investigated the root tissue types in which both warm- and cold-induced FRI condensation occurred. Under warm temperatures, we observed the presence of FRI condensates in the division and growth zones of a subset of emerging and young lateral roots but not in primary roots (Fig. 6C, Nplants=8/29 lateral roots, Nplants=0/34 primary roots of 6-13 DAG). This was different to what occurred under cold temperatures, in which we found FRI condensing in all root types, including the primary roots (Supplemental Fig. 7). Interestingly, the occurrence of warm-induced FRI condensates within a given lateral root cell was highly variable among lateral roots, with detectable FRI-mScarletI condensates found in anywhere from 30% to 94% of the epidermal and cortical cells within the growth zone (Supplemental Fig. 7). The seemingly stochastic patterns of nuclei containing FRI condensates among and within lateral meristems resemble the previously described phenomenon of transcriptional and translational “bursting,” which has been associated with epigenetic and phenotypic diversity in response to environmental changes^17,29,30^.

Previous research has highlighted the influence of plant tissue types on the state of auxin response factor (ARF) biomolecular condensates^31^. The observation of co-condensation between SUF4 and FRI in lateral roots, while absent in primary roots of 6-13 days after germination plants, suggests the potential for a tissue-specific functional relationship between these proteins. If this is true, we might observe a tissue-specific change in mScarletI-FRI condensate behavior in response to changes in SUF4 condensation. To test this hypothesis, we introduced FRI-mScarletI into *suf4* null, 12S SUF4, and 5S SUF4 variant plants. At room temperatures, we observed the formation of FRI-mScarletI condensates in about 10%-25% of primary roots of *suf4* null and 12S SUF4 plants, respectively (Fig. 6C, Table. 1; suf4 4/41, 12S SUF4 4/5), but no FRI-mScarletI condensates were observed in the primary roots of rescue SUF4 (0/34) or 5S SUF4 lines (0/19). FRI-mScarletI condensates were observed in a portion of lateral roots in all lines. Although the number of mGFP-SUF4 condensates varies between rescue SUF4 and 5S SUF4, both genotypes have the ability to form condensates, whereas 12S and *suf4* (*FRI*) do not. Thus, our data indicate an inverse relationship exists between SUF4 and FRI condensates in the primary root. Furthermore, they suggest that FRI is capable of forming at least two distinct types of condensates that vary by tissue type: one type that assembles with SUF4, likely functioning to help establish *FLC* transcription in newly formed tissue, and another type where the proteins possibly act antagonistically.

**Table 1.**
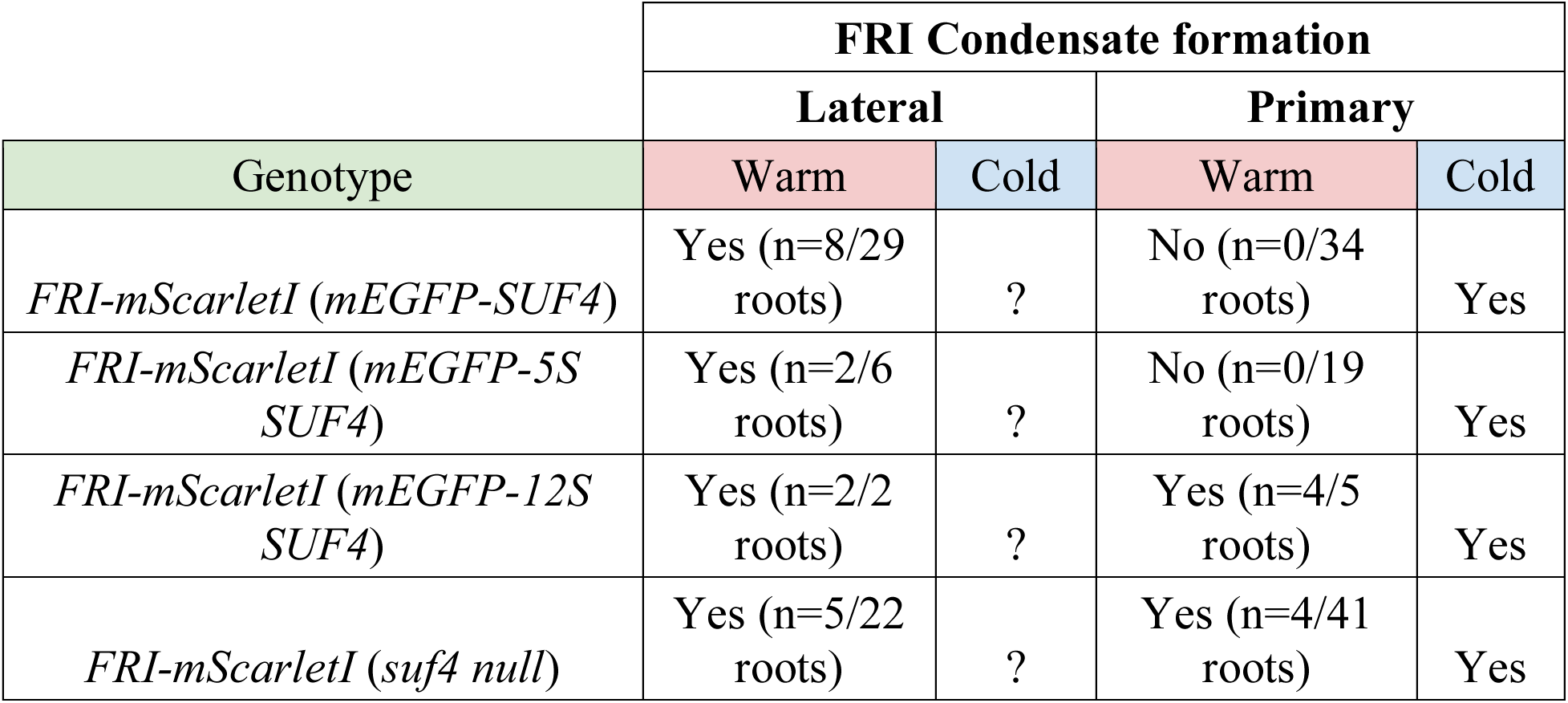
FRI-mScarletI condensate formation in different root types within SUF4 variants.

### SUF4 colocalizes with ELF7 in nuclear condensates at room temperature

While FRI facilitates transcriptional activation of *FLC* by establishing a permissive chromatin environment, other mechanisms have been shown to be involved in the maintenance of *FLC* expression, such as transcriptional regulation associated with histone H3 lysine 4 (H3K4) modifications ^32,33^. In light of this, we sought to investigate whether SUF4 condensates might serve as a hub for multiple proteins involved in the transcriptional regulation of *FLC* by fluorescently tagging and expressing EARLY FLOWERING 7 (ELF7), a known member of the PAF1 complex. We introduced mTFP-ELF7 expressed from its own upstream sequence (pELF7::mTFP-ELF7) into rescue lines expressing mEGFP-SUF4 and mEGFP-SUF4/FRI-mScarletI. ELF7 was specifically chosen due to its previous genetic interaction with SUF4 and its role in establishing H3K4 activation marks on *FLC*, as well as its physical co-localization with FRI in nuclear condensates under normal temperature conditions^26,32,33^. Within both transgenic lines, we observed mTFP-ELF7 to be diffuse within the nucleoplasm of cortical and epidermal cells within the transition zone of both lateral and primary roots. In a subset of these cells, mTFP-ELF7 formed 1-2 prominent nuclear condensates and occasionally multiple smaller condensates (Fig. 5B).

As the introduction of mTFP-ELF7 did not appear to induce silencing, nor alter condensate behavior in either the rescue mEGFP-SUF4 or mEGFP-SUF4/FRI-mScarletI lines, we performed colocalization analysis using lines expressing all three transgenes (Fig. 5B). While mEGFP-SUF4 formed more condensates compared to ELF7 and FRI in nuclei analyzed (nSUF4condensates= 414, nELF7condensates= 53, nFRIcondensates= 34), the presence of an ELF7 condensate almost always coincided with a SUF4 condensate (47 out of 53 cases). Further, out of these 47 ELF7-SUF4-containing condensates, we observed the additional presence of a FRI condensate in 8 cases (17%). The quantitative correlation between SUF4 condensation and flowering time, coupled with its ability to co-localize with both FRI and ELF7 in 1-2 condensates, supports the notion that SUF4 condensation likely functions in regulating the transcriptional activity of *FLC* under different temperature conditions (Fig. 7).

**Figure 7:**
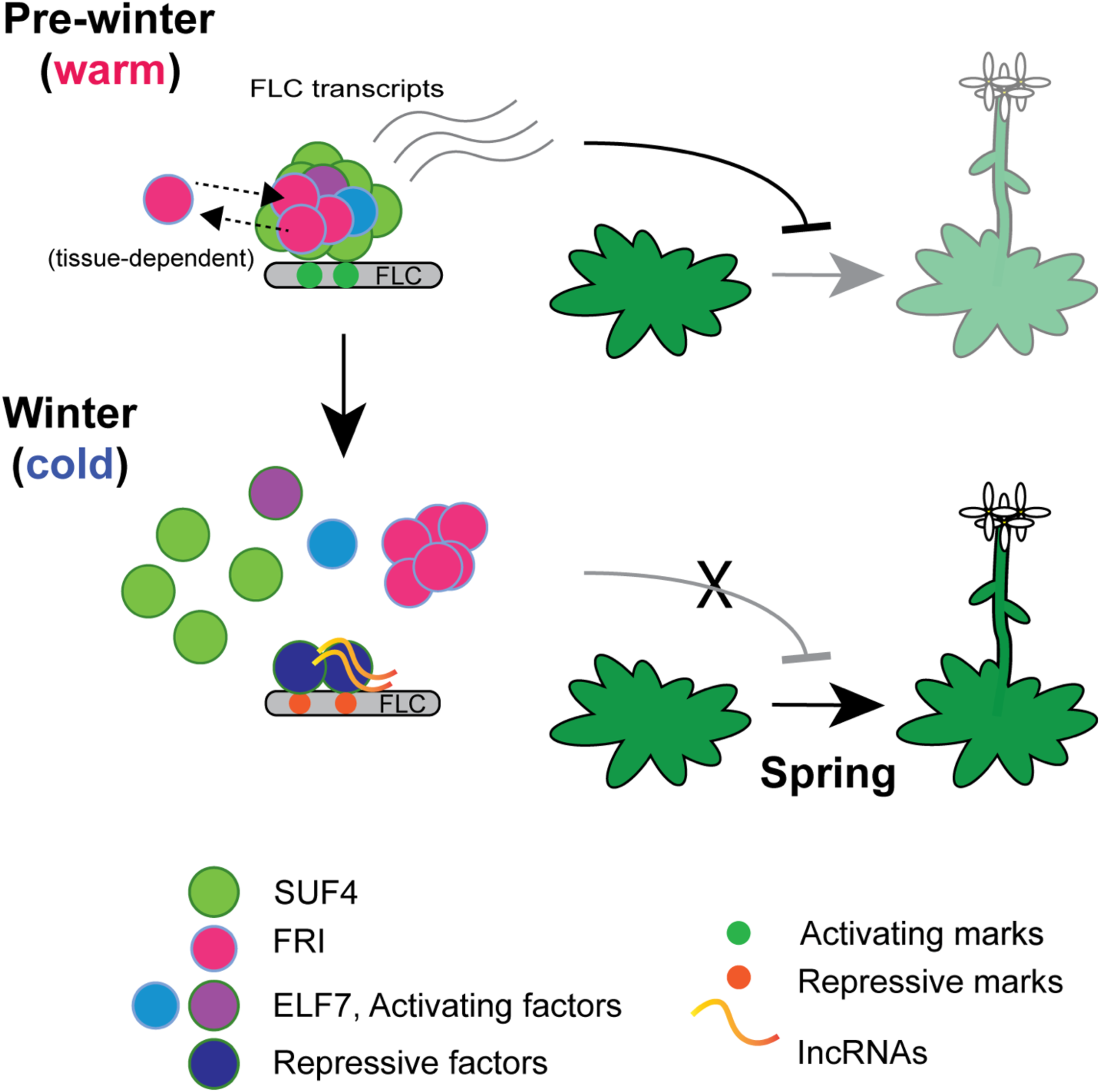
Proposed Model for Flowering-Time Regulation. A proposed model for the regulation of flowering time mediated by SUF4 biomolecular condensates. SUF4 (green) biomolecular condensates function to concentrate and/or recruit members of the vernalization complex, including FRI (in red), ELF7 (in blue), and other associated factors (in purple) to the *FLC* gene locus. Certain members, such as FRI, exhibit transient localization within these condensates dependent on cell state. Condensate localization ultimately leads to the transcriptional activation of the *FLC* gene, resulting in the delay in flowering time. In response to cold temperatures, SUF4 condensates undergo disassembly. This disassembly may facilitate the recruitment of a new group of chromatin remodelers and the involvement of lncRNAs, both of which work together to transcriptionally silence the *FLC* gene. Transcriptional silencing of FLC is a necessary step for plants to undergo the floral transition and flower in the subsequent spring season.

## DISCUSSION

In this study, we have discovered that the IDP, SUF4, undergoes thermally-regulated condensation both *in vitro* and *in planta*. These findings, along with the demonstrated ability of SUF4 sequence alterations to progressively tune phase separation, *FLC* transcription, and predictable flowering time outcomes, and the evidence that SUF4 condenses with *FLC* transcriptional regulators under appropriate physiological conditions, provide strong support for the hypothesis that SUF4 phase separation may serve as a physical mechanism for plant temperature perception. Our work emphasizes the biological importance of protein phase separation in a multicellular organism by establishing links between environmentally-sensitive phase separation, gene expression, development and physiology.

Plant cells require an ability to filter out noise and interpret meaningful signals from their environment to make developmental decisions. The conformational promiscuity and multivalency of IDPs enables them to engage in a variety of specific yet transient interactions under defined environmental conditions. IDPs are enriched within plant proteomes, with many participating in plant regulatory pathways and as integrators of environmental inputs^34–36^. The phase separation states of such IDPs and client proteins could serve as an effective mechanism for distinguishing real signals from noise within plant cells, thereby enabling more efficient and effective developmental outcomes. Plants, along with other organisms, use epigenetic factors, such as changes in chromatin/histone state, signaling pathways, and RNA modifications to promote environmental-driven developmental changes. However, these changes often require integration over a significant period of time in order to induce a physiologically-relevant response. Pairing IDP sensing with slower-acting epigenetic mechanisms may allow plants to better distinguish genuine signals from noise by facilitating the promotion or maintenance of molecular memory under persistent environmental cues. In the case of SUF4, we found that changes in temperature cause it to phase transition between condensed and uncondensed forms. Moreover, manipulating the aromatic/hydrophobic properties of SUF4 proved to be sufficient to modulate its material characteristics and tune the onset of flowering. Given SUF4’s known interaction with *FLC*-related chromatin remodelers and our cell biological evidence that at least one of these co-condenses with SUF4 *in vivo*, these findings suggest the possibility that SUF4 condensates could also associate with and alter the activity of epigenetic factors to modulate *FLC* transcription.

Extensive investigations have shed light on the intricate regulation of flowering time in *Arabidopsis*, with a specific emphasis on the transcriptional modulation of *FLC*^1^. Among the key players in this regulatory network is FRI, an interacting partner of SUF4^2,36^. Equipped with two coil-coil domains, FRI is proposed to act as a scaffold protein, facilitating interactions with various *FLC* transcriptional regulators and chromatin remodelers. Nevertheless, FRI itself does not directly interact with the *FLC* gene locus. Instead, SUF4 serves as a mediator of this interaction by physically binding both *FLC* and FRI, along with many other members associated with the FRI complex ^3,37^. Through both condensation and its ability to recognize the *FLC* promoter region, SUF4 has the potential to enhance its own activity by raising its local concentration, and also possibly by recruiting and concentrating transcriptional activators (e.g. previously reported SUF4 interactors FLX and TAF14), and chromatin remodelers (e.g. ELF7, SWC6, and RVB1) at the *FLC* promoter. Meanwhile, we hypothesize that FRI transiently interacts with SUF4, contributing to the establishment of transcriptional regulation of *FLC* in newly formed tissues such as lateral roots. Newly developed and emerging molecular images and technologies, such as those that enable the visualization and quantitation of molecular events at a single genetic locus (e.g., endogenous fluorescent tagging of gene loci via dCAS9/CRISPR), will aid in further investigating the functional dynamics that govern these interactions at the *FLC* gene locus.

The coacervation of prion-like proteins (PrLPs), such as FLL2 and FCA, has recently emerged as a potential regulatory mechanism in flowering time ^38^. However, it remains uncertain whether PrLP coacervation is thermally-driven, or even contributes to the post-transcriptional regulation of *FLC*. In this study, we demonstrate that a distinct proline-rich IDP undergoes LCST condensation behavior, which likely serves as a means to modulate flowering. This behavior aligns with the LCST observations of synthetic elastin-like polypeptides and Pab1 ^22,39–42^, which also possess a high proline content and repetitive hydrophobic features. While plants carry numerous proline-rich proteins and hybrid proline-rich proteins that are involved with abiotic- and biotic-associated pathways, the thermo- induced coacervation behavior of these proteins remains underexplored. By unraveling the role of proline-rich IDPs in phase separation and its relevance to flowering time regulation, our study extends our understanding of proline-rich IDPs in biological processes and sheds light on their potential participation in eukaryotic thermo-responses through phase separation phenomena. Moreover, our findings provide a strong basis for exploring the role of phase separation and environmental sensing, providing a valuable foundation and new insights into how we may be able to engineer plants and other organisms with tailored environmental responses.

Despite IDPs being often poorly conserved ^43,44^, SUF4 shares substantial homology with the animal protein BuGZ. In these organisms, BuGZ plays essential roles in both mitotic and transcriptional regulation by interacting with various cellular machineries, including the spindle assembly checkpoint protein (SAC) Bub3, the spliceosome, and epigenetic factors^10–12^. While SUF4’s primary function in plants revolves around regulating flowering time, recent research has unveiled its potential involvement in analogous mitotic and transcriptional regulatory processes ^45–47^. These newfound roles raise a series of questions, including whether SUF4’s phase separation has evolved distinct functionality specific to plant flowering or if it serves as a universal mechanism within eukaryotic systems. Interestingly, animal BuGZ has been observed to undergo phase separation *in vitro*, and its role in phase separation is hypothesized to contribute to mitotic spindle formation. Therefore, studying SUF4 may not only advance our comprehension of how environmentally-induced phase separation impacts plant development and physiology but also hold significant promise in deepening our understanding of the broader evolutionary significance and functional implications associated with such environmentally-induced phase separation.

## MATERIALS AND METHODS

### Plant Material, accessions, and growth conditions

All plants used in this study were grown under controlled conditions in growth rooms with a light intensity of approximately 180 µmol m−2 s−1 at a temperature of 21°C. Plants were either exposed to long day conditions (16 hours of light followed by 8 hours of darkness).

*Col (FRI-SF2)* ^48^ was used as the wild-type accession for all genotypes analyzed in this study. These lines are characterized as winter annuals and exhibit a delayed flowering phenotype.

*suf4 (FRI-SF2)* plants ^32^ were used as the mutant accession for this study. These plants carry an ∼6kb deletion that spans the N-terminus of SUF4 (deleted region: exons 1,2,3 and part of intron 3), resulting in a null mutation that causes an early flowering phenotype.

| Transgenic lines generated in this study |
| --- |
| pSUF4::mEGFP-SUF4 in *suf4 (FRI-SF2)* |
| pSUF4::mEGFP-5SSUF4 in *suf4 (FRI-SF2)* |
| pSUF4::mEGFP-12SUF4 in *suf4 (FRI-SF2)* |
| pFRI::FRI-mScarletI; pSUF4::mEGFP-SUF4 in *suf4 (FRI-SF2)* |
| pFRI::FRI-mScarletI; pSUF4::mEGFP-5SSUF4 in *suf4 (FRI-SF2)* |
| pFRI::FRI-mScarletI; pSUF4::mEGFP-12SSUF4 in *suf4 (FRI-SF2)* |
| pFRI::FRI-mScarletI; pSUF4::mEGFP-30ISUF4 in *suf4 (FRI-SF2)* |
| pELF7::mTFP-ELF7; pSUF4::mEGFP-SUF4 in *suf4 (FRI-SF2)* |
| pELF7::mTFP-ELF7; pFRI::FRI-mScarletI; pSUF4::mEGFP-SUF4 in *suf4 (FRI-SF2)* |
| U2B"-mRFP(pROK2); pSUF4::mEGFP-SUF4 in *suf4 (FRI-SF2)* |

### Flowering time quantification

To assess flowering time in SUF4 variants, seeds of each genotype were individually sown in Pro Mix BX soil and stratified at 4°C for two days. Then, planted pots were transferred to either short day or long day chambers. The total number of true rosette leaves was recorded every 3-5 days until plants bolted. To minimize position effects, all genotypes were spatially mixed within their respective short- or long day growth chambers.

### Cloning schemes

#### pHM11 (rescue SUF4)

To create pHM11 (pSUF4::6HIS-mEGFP-SUF4), a 2,981 fragment containing the genomic SUF4 sequence was amplified using oHM3 (TTA GGA TCC GGT GGA GGC GGT TCA GGC GGA GGT GGC TCT GGC GGT GGC GGA TCG ATG GGT AAG AAG AAG AAG AGA), a forward primer with an added BamHI site and a 3XGGGS linker, and oHM4 (GGC CTC GAG CTA AAA CGC CAT CCG CCC AGC AA), a reverse primer with a 3’ XhoI site. This fragment was then cloned into pGEM-T easy to generate pHM1 (genSUF4). Additionally, the mEGFP gene was amplified from addgene’s commercially available mEGFP plasmid (#18696) using oHM1 (ATA AGA TCT AGT GAG CAA GGG CGA GGA G), a primer with a BglII 5’ site, and oHM2 (ATG GAT CCC TTG TAC AGC TCG TCC ATG CC), a primer with a 3’ BamHI site. The resulting fragment was cloned into pGEM-T easy to create pHM2 (mEGFP). Subsequently, pHM2 was digested with BglII and BamHI, and then inserted into the pET-30a(+) vector, resulting in pHM3 (6HIS-mEGFP). Plasmids pHM1 and pHM3 were then cleaved with BamHI and XhoI and ligated, yielding pHM4 (6HIS-mEGFP-SUF4). A 1059 bp fragment upstream of the SUF4 coding region was PCR amplified using oHM28 (ATC TTT AAC TTT TTT GTT GTT GCA) and oHM24 (GAT GGC GCG CCC ACT AAA TCT TTC TCC TA), a primer with an ASCI site, and inserted into the pGEM-T easy vector, generating pHM5 (proSUF4). Additionally, a 332 bp region downstream of the SUF4 coding region was PCR amplified using oHM25 (GAT GGC GCG CCC TCG AGA ATT TGG CAC CAA ACC AA), a primer with both ASCI XHOI restriction sites, and oHM17 (gaacaacatggtgcaaacatt), and cloned into pGEM-T easy to produce pHM6 (termSUF4). Subsequently, pHM5 was digested with SacII and AscI, and the resulting fragment was cloned into pHM6 to create pHM7 (proSUF4::AscI-XhoI-3’UTR). pHM4 (6HIS-mEGFP-SUF4) was reverse amplified to add a kozak consensus sequence using primers ohm45 (TTCACCAAAATGCACCATCATCATCATCAt) and oHM46 (GGcgcgCCCACTAAATCTTTCTCC), and then digested with AscI and XhoI for cloning into pHM7, resulting in pHM10 (proSUF4::6HIS-mEGFP-SUF4-3’UTR).

For downstream biochemical assays requiring removal of mEGFP, an enterokinase sequence was introduced before the SUF4 coding region using primers oHM35 (CGATCCGCCACCGCCAGAGCC) and oHM36 (gacgacgacgacaagATGGGTAAGAAGAAGAAGAGA), generating pHM10_entero (proSUF4::6HIS-mEGFP-entero-SUF4-3’UTR). Subsequently, pHM10_entero was cloned into pBIN30 via the NotI restriction site to make pHM11 (proSUF4::6HIS-mEGFP-entero-SUF4-3’UTR).

#### pHM32 (30I SUF4)

To create pHM32, the last 1969 bp of SUF4’s disordered region was synthesized (GenScript, USA) with base pair substitutions that replaced all 30 alanine residues with isoleucine residues. Gibson assembly was used to fuse this fragment with the remaining 1012 bp genomic sequence of SUF4 to reassemble the complete gene. Once the 30I SUF4 fragment was assembled, pHM10_entero and the 30I fragment were cut with BamHI and XhoI and ligated together to make pHM31 (pSUF4::mEGFP-30ISUF4). Then, pHM31 was cut with Not1 and cloned into the binary vector pBIN30 (Basta resistant) to make pHM32 (pSUF4::mEGFP-30ISUF4).

#### pHM14 (12S SUF4)

To create pHM14, the last 1969 bp of SUF4’s disordered region was synthesized (GenScript, USA) with base pair substitutions that replaced 12 of the 14 phenylalanine and lysine residues with serines. Gibson assembly was used to fuse this fragment with the remaining 1012 bp genomic sequence of SUF4 to reassemble the complete gene. Once the 12S SUF4 fragment was assembled, pHm10_entero and the 12S fragment were cut with BamHI and XhoI and ligated together to make pHM12 (pSUF4::mEGFP-12SSUF4). Then, pHM12 was cut with Not1 and cloned into the binary vector pBIN30 (Basta resistant) to make pHM14 (pSUF4::mEGFP-12SSUF4).

#### pHM5S (5S SUF4)

To create pHM5S, the last 1969 bp of SUF4’s disordered region was synthesized (GenScript, USA) with base pair substitutions that replaced the last 5 phenylalanine and lysine residues with serines. Gibson assembly was used to fuse the remaining 1012 bp genomic sequence of SUF4 to reassemble the complete gene. Once the 5S SUF4 fragment was assembled, pHm10_entero and the 5S fragment were cut with BamHI and XhoI, and ligated together to make pHM13 (pSUF4::mEGFP-5SSUF4). Then, pHM13 was cut with Not1 and cloned into the binary vector pBIN30 (Basta resistant) to make the final vector pHM5S (pSUF4::mEGFP-5SSUF4).

#### pFRI::FRI-mScarletI

pFRI::FRI-mScarletI was custom-synthesized by GenScript and consists of the following elements: The FRI 1243 bp upstream promoter region with an added Kozak consensus sequence (CACCAAA), directly followed by the 1351 bp FRI (SF2) genomic region, 3XGGGS linker (GGT GGA GGC GGT TCA GGC GGA GGT GGC TCT GGC GGT GGC GGA TCG), mScarletI sequence derived from Addgene 98839, and FRI’s putative 169 bp 3’UTR. Both promoter and 3’UTR sequences were obtained from TAIR Sequence Viewer, whereas the genomic sequence was provided by Sang Yeol Kim. The aggregated synthesized DNA was flanked with NotI cut sites and subsequently inserted into the pBIN40 (Hyg resistant) binary vector through NotI digestion and ligation.

#### pELF7::mTFP-ELF7

pELF7::mTFP-ELF7 was custom synthesized by GenScript, USA and comprises the following components: The ELF7 405 bp upstream promoter region with an added Kozak consensus sequence (CACCAAA), directly followed by the mTFP sequence (Addgene 55503), 3XGGGS linker (GGT GGA GGC GGT TCA GGC GGA GGT GGC TCT GGC GGT GGC GGA TCG), the 3228 bp ELF7 genomic region (TAIR), and the ELF7 putative 1791 bp 3’ UTR. All promoter, genomic, and 3’UTR sequences were obtained from TAIR Sequence Viewer. The synthesized DNA was flanked by NotI sites, which were used for subsequent insertion into the pBIN40 (Hyg resistant) binary vector.

#### U2B”-mRFP (pROK2)

U2B” -mRFP (pROK2) was previously created by Alie Pendle and Peter Shaw and commercially available for order via ABRC (https://abrc.osu.edu/stocks/number/CD3-795056).

### Live Cell Microscopy and Fluorescence Recovery After Photobleaching (FRAP)

Live-cell imaging was performed on roots of 6-7 day old seedlings using either point scanning or a spinning disk scanning confocal microscopy. Point-scanning was performed on a Leica SP8 microscope equipped with HyD detectors, a white light supercontinuum laser, and AOBS (Acousto-Optical Beam Splitter) control of shuttering, excitation and emission bands. All images were acquired with a Leica 60x 1.3 N.A Plan-Apo glycerin immersion objective. Leica Lightning deconvolution was used to improve signal to background for all point-scanning images. Spinning disk confocal microscopy was performed either with a CSU-W110 confocal head (Yokogawa) combined with a MESA Field-flattening device (Intelligent Imaging Innovations) mounted on an Olympus IX83 microscope equipped with a 100x Plan-Apo 1.35 NA silicon oil immersion objective, or a fully automated Leica DMI6000B microscope equipped with a CSU-X spinning disk confocal head (Yokogawa Corp.). Each sample was excited using 488nm for GFP (Coherent cube laser), 561nm for mScarletI (Coherent cube laser), or 442nm for mTFP (Coherent cube laser). Emitted light was collected with a 525/50 nm filter for GFP, 605/64 nm filter for mScarletI and a 482/35 nm filter for mTFP (all filters from Semrock).

FRAP experiments were conducted using the CSU-X spinning disk microscope described above, with aid of a Vector laser scanning unit (Intelligent Imaging Innovations) attached to the right camera port to focus and steer laser energy on the image plane. The camera port was switched from imaging to bleaching paths via software control (SlideBook, Intelligent Imaging Innovations) to alternate between imaging and FRAP light paths. mEGFP-SUF4 was excited using 3.5 mW of light energy (as measured at the end of the laser fiber entering the confocal head) from a solid state 488nm laser (Coherent Technologies). Emission light was collected using a 488nm dichroic mirror and 500-550nm emission filter (Semrock). Single-plane Images were collected every 200ms using a Leica 100x Plan-Apochromat 100x 1.40 NA oil immersion objective and FRAP was performed within a 1.094 micron^2^ region of interest centered on each assessed condensate spot using 5mW of light energy as measured at the end of the fiber entering the Vector unit. The focal spot was moved by galvometers to bleach a grid of spots separated by 20 nm within the FRAP region. Seven frames were collected before bleaching and 143 frames were collected after bleaching (1.75 seconds to conduct bleaching, 0.2 second delay to first post-bleach acquisition). Once all images were acquired, they were aligned in ImageJ ^49^ using the MultiStackReg plugin ^50^. The Trackmate plugin ^51^ was used to segment mEGFP-SUF4 spots for measurement of both the mean fluorescent intensity and mobility of condensates over time (DoG detector, estimated object diameter 0.6µm, sub-pixel localization, simple LAP tracker: Linking max distance 1µm, Gap-closing distance 1µm, and Gap-closing max frame gap 1). Afterwards, easyFRAP ^52^ (https://easyfrap.vmnet.upatras.gr/) was used to calculate double normalized (initial intensity, photobleaching) FRAP values among the different variants, with each curve undergoing a double exponential curve fit (Normalization method: Double, Exponential Equation: Double, Data collected: T-Half, Mobile Fraction, R-Squared).

### Live-cell imaging during temperature change

Temperature change experiments were performed on the Leica SP8 microscope. Seedlings were placed in a 4C growth chamber for 24 hours. Then, they were mounted in ice water and moved onto the microscope stage, where they underwent continuous imaging for 10 minutes. As a control, room temperature seedlings were mounted in room temperature water and continuously imaged using the same amount of time.

### Colocalization analysis

Co-localization of mEGFP-SUF4 condensates with FRI-mScarletI, U2B”, and/or mTFP-ELF7 was investigated by collecting multichannel 3D datasets on the spinning disk microscope with the CSU-W1 confocal head described above. Images were acquired with a 100x Plan-Apo 1.35 NA silicon oil immersion objective at 0.3µm intervals on the z-axis. The Spots function and Matlab plugin for colocalization analysis in Imaris software (RRID:SCR_007370, URL:http://www.bitplane.com/imaris/imaris) were used to segment condensate signal and assess co-localization. Spot diameters of 0.5µm were used to detect and determine the locations of mEGFP-SUF4, whereas FRI and U2B” condensates were detected with a spot diameter of 0.7µm. These diameters values were determined by assessing the diameter that worked best for segmentation of condensate signal using the Imaris Spots function and were largely indicative of the typical condensate diameters. Spots were considered co-localized if the centroids of the spots (determined at a sub-pixel scale), were closer than the radius of the smaller Spot size used to detect and segment mEGFP-SUF4 signal (0.25 µm).

### Puncta quantification

To quantify SUF4 nuclear condensates, we auto-cropped individual nuclei from 3D stacks using the NucleusJ2.0 auto crop function in Fiji. Subsequently, the cropped images were imported into LabView, where a custom signal algorithm was applied to normalize the signal loss due to light scattering and spherical aberration from the top to the bottom of the z-stacks. To create the dataset used for normalization, we isolated the diffuse EGFP-SUF4 signal within the nucleoplasm away from the brighter SUF4 puncta via a set of binary masks, one mask for total SUF4 signal above background and second mask for the puncta. The threshold for the total mask was set by eye, and that for the puncta was created to include only pixels falling between 0.02% and 20% below the maximum pixel value in the section. For each section, the puncta signal was then subtracted from the total signal to obtain the nucleoplasm signal, and the mean nucleoplasm value per pixel calculated. The nucleoplasm signal was roughly linear to the focal depth once the focal plane was within the nuclear volume, enabling us to model signal loss with a linear fit to the mean intensity in these focal sections. The pixels in planes conjugate the nuclear volume were normalized to the inverse of the fit line. The code for TIFF file reading was shared online by Brian Hoover(<u>forums.ni.com</u>), the code for TIFF multipage saving was shared online by Nico (<u>forums.ni.com</u>) and used with modifications.

### Statistical analysis

For the analysis of flowering time and condensate numbers, a nonparametric Mann-Whitney U test was used to test for statistically significant differences among sets of measurements made in different genotypes. For qPCR, FRAP and motility results, a Student’s two-tailed T-test for independent means was used. To assess the significance of the observed co-localization frequency for FRI-mScarletI and mEGFP-SUF4 condensates, we compared this frequency to that observed for a control protein U2B”-mRFP, which forms 1-2 bodies per nucleus of similar sizes to those observed for FRI-mScarletI. This experiment assumes that all three proteins are localized to the nucleosol and allows for a statistical comparison without knowing the spatial structure and volume of the nucleosol. A two-sample (binomial) test of proportions was used to analyze the results.

### QPCR

qPCR was employed to assess the transcript levels of both *SUF4* and *FLC*. To achieve this, total RNA was extracted from six-day-old seedlings (approximately 20 seedlings per genotype) using the NucleoSpin RNA plant kit (Macherey-Nagel CO.). Subsequently, 1 microgram of total RNA was reverse transcribed using the QuantiTect Reverse Transcription kit (Qiagen). Real-time RT-PCR was performed on 1:10 diluted cDNA using the SensiFAST SYBR No-ROX Kit (Bioline). Three biological and technical replicates were analyzed per genotype, and ROC1 (AT4G38740) was used as a reference gene to normalize gene expression. The reactions were carried out with an initial denaturation step at 95°C for 2 min, followed by 45 cycles of 95°C for 5 s, 60°C for 10 s, and a final step at 72°C for 5 s.

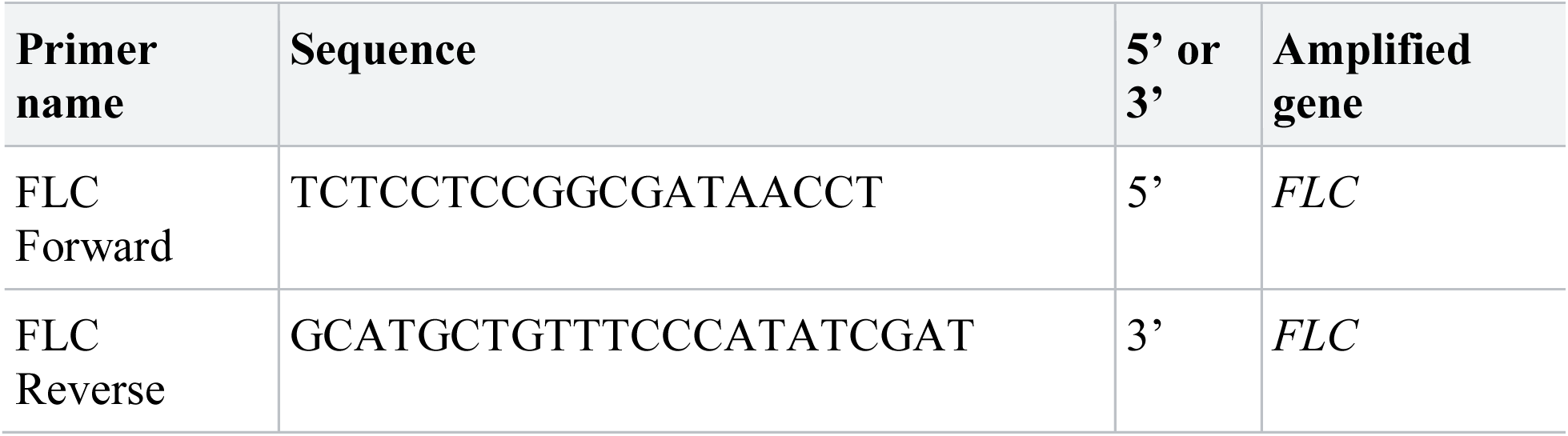
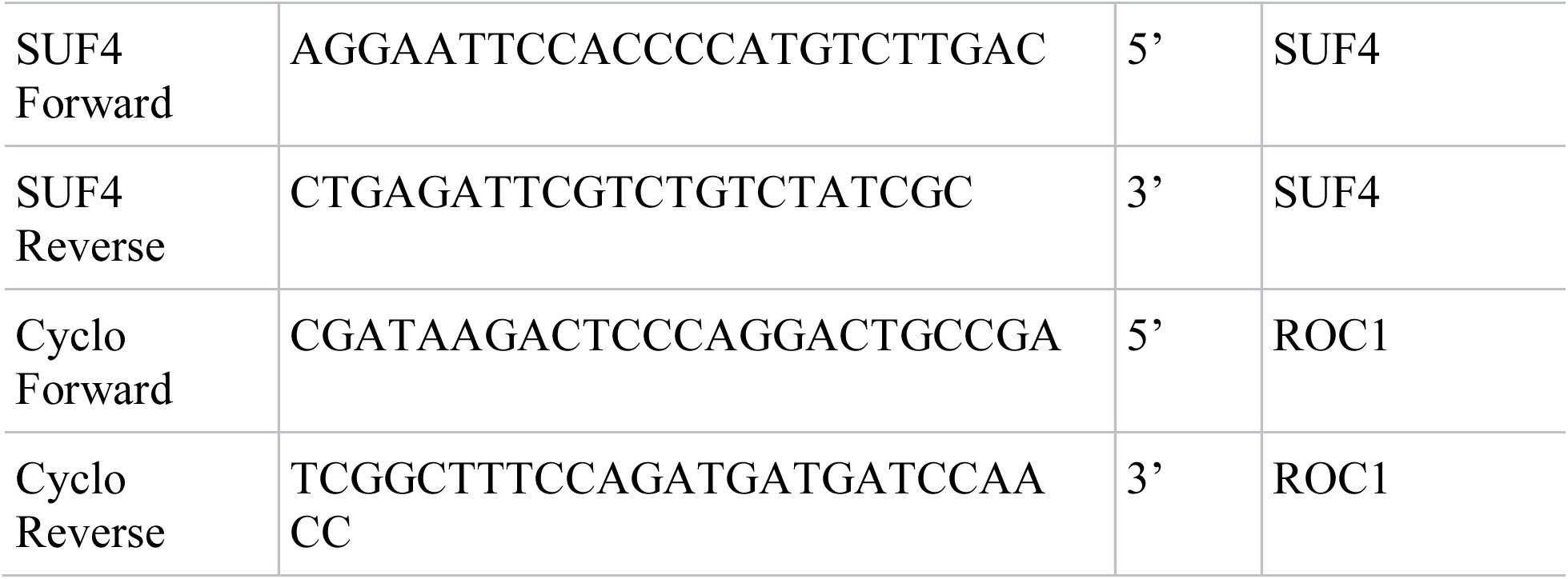

### Expression of His-mVenus-SUF4 in Sf9 cells

Cloning and expression of His-mVenus-SUF4 was based on the Bac-to-Bac Baculovirus Expression system (ThermoFisher). The control pFastBac-mVenus and pFastBac-mVenus-SUF4 isoform 1 plasmids were obtained from Dr. Alex Chin (*15*). In these vectors, mVenus was cloned solely or with an open reading frame of SUF4 isoform 1 (NP_564369.1; 367 amino acid) in the modified pFastBac vector (ThermoFisher). Site-directed mutagenesis was carried out to create the 5S and 12S mutants using PrimeSTAR Max DNA polymerase (Takara-bio). All plasmids were sequenced and verified prior to the transformation of DH10Bac. Bacmid preparation, transfection and infection of Sf9 cells were performed according to the manufacturer’s instruction. Large scale protein expression was carried out by infecting and culturing the Sf9 cells with P3 viruses for 2 days. Cells were centrifuged and either used immediately for protein purification or frozen and stored at -80°C until use. Purification of His-mVenus-SUF4 fusion protein from Sf9 cells. His-mVenus-SUF4 proteins were purified with IMAC followed by gel filtration as published previously with some modifications. All procedures were performed at 4°C. Proteins were extracted by sonicating the cells in lysis buffer [50 mM Tris, 500 mM NaCl, 20 mM imidazole, 5 mM TCEP (SIGMA, C4706), 5% glycerol, protease inhibitor cocktail (SIGMA, S8830) pH 7.44] supplemented with benzonase nuclease (SIGMA, E1014). The lysate was cleared by centrifugation at 20,000 x g, for 20 min at 4°C, filtered with 0.45 µm PVDF membrane filter (Millipore, SLHV033RS) and loaded onto Ni-NTA columns (QIAGEN, 30230). After washing with the lysis buffer, proteins were eluted with 500 mM imidazole (lysis buffer supplemented with 480 mM imidazole). The eluate was concentrated using a centrifugal filter device, Amicon Ultra-15 (Millipore, UFC901024) and loaded to a gel filtration column Superdex 200 Increase 10/300 GL operated with AKTA Pure 25L (GE Healthcare). Gel filtration was performed in the Size Exclusion Chromatography (SEC) buffer (20 mM Hepes, 225 mM NaCl, 1 mM TCEP pH 7.4) with a constant flow rate of 0.4 ml/min. Typically, His-mVenus-SUF4 and His-mVenus proteins were eluted at 11.0 and 15.5-ml after the protein loading, respectively. Obtained fractions were analyzed with SDS-PAGE, combined and concentrated. Protein concentration was determined with Bradford protein assay kit (Thermo, 23200) using BSA as a standard.

### Turbidity assay

His-mVenus or His-mVenus-SUF4 proteins were thawed and ultracentrifuged to remove denatured proteins. After protein concentration was determined, the protein sample was diluted to 50, 25, 12.5, 6.25 and 3.13 µM in SEC buffer. Each sample was mixed well with equal volume of turbidity assay buffer (20 mM Hepes, 75 mM NaCl, pH 7.4) to make a 250 µl reaction mix and transferred into a quartz cuvette (Starna Cells, 28F-Q-10). Temperature-dependent phase transition was detected by measuring the optical scattering at 800 nm while increasing the temperature from 4 to 60°C (with a ramping rate of 0.5°C/min) using a spectrophotometer Cary 300UV-VIS (Agilent Technologies). Nitrogen gas purging was used to prevent the moisture from condensing on the cuvette surface. Three replicates were performed for each sample.

### Droplet formation assay

Five microliter of 50 µM proteins prepared as described above were mixed with 5 µl of turbidity assay buffer on ice and incubated for 5 min at 37°C. Each 3 µl of protein reaction mix was loaded into a flow-chamber and immediately observed with a confocal microscope SP5 (Leica) equipped with a 60x oil objective (NA 1.42). Imaging was performed with both DIC and fluorescence (excitation at 514 nm, emission 530 nm).

## Supplementary Materials

**Supplemental Figure 1.**
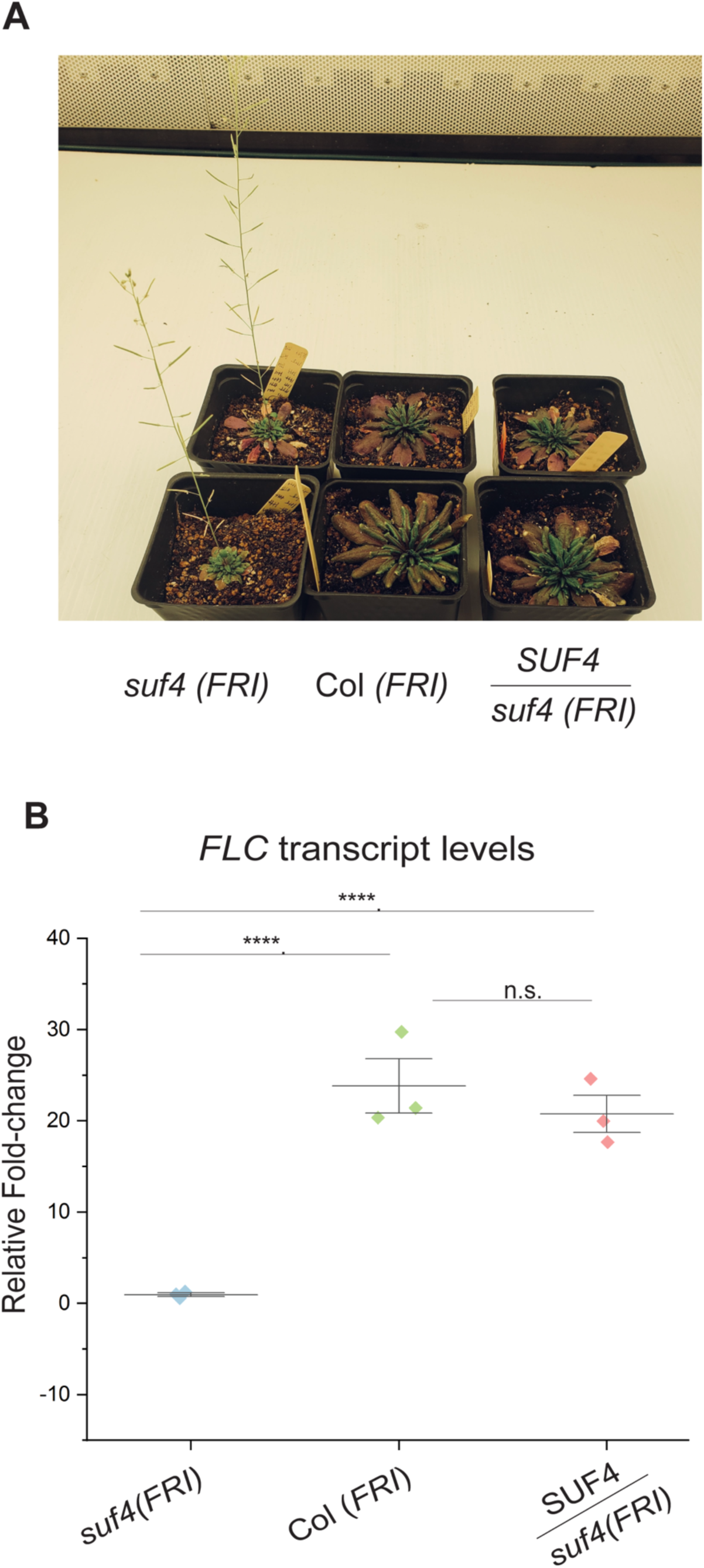
Expression of the pSUF4::mEGFP-SUF4 transgene rescues the *suf4* mutant phenotype **A)** Image of plants demonstrating the ability of the pSUF4::mEGFP-SUF4 transgene to rescue the *SUF4* flowering time phenotype. From left to right: *suf4 (FRI)* mutant plants displaying an early flowering time phenotype, Col *(FRI)* plants displaying a WT delayed flowering time phenotype, SUF4/*suf4 (FRI)* displaying a rescued WT delayed phenotype. **B)** Relative abundance of *FLC* transcripts in *suf4* mutant, Col *(FRI)*, and *SUF4 (FRI)* shown in A). Rescue SUF4 shows FLC transcript levels similar to WT. These data are reshown in Figure 2E. Statistical analysis conducted using a Student’s t-test. Error bars represent the standard deviation. Asterisks denote the p-values* <0.01, ** <0.001, ***<0.001, and ****<0.0001.

**Supplemental Figure 2.**
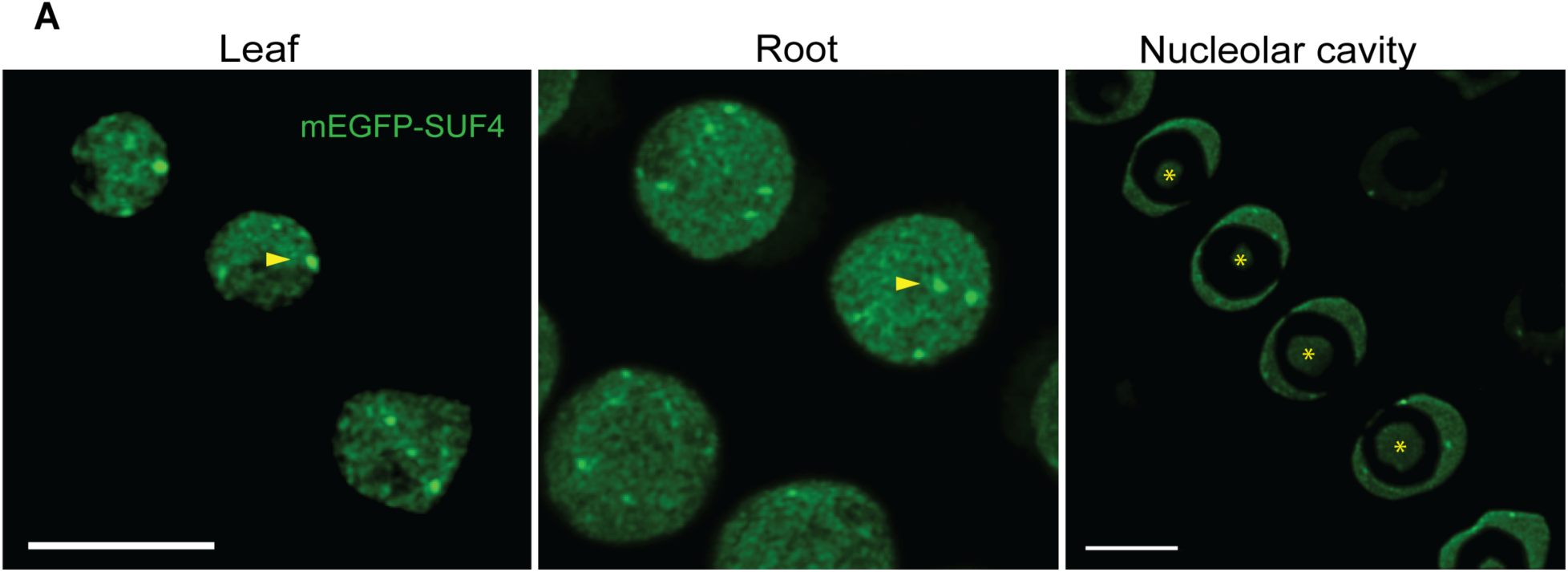
At room temperatures, SUF4 forms biomolecular condensates in shoot cells, primary root cell, and localizes in the nucleolar cavity. Arrows point to nuclear condensates whereas stars mark the nucleolar cavity. Leaf and Root images are 3D rendered Z-stacks whereas the nucleolar cavity image is a slice from a 3D stack. Scalebar: 10 µm

**Supplemental Figure 3.**
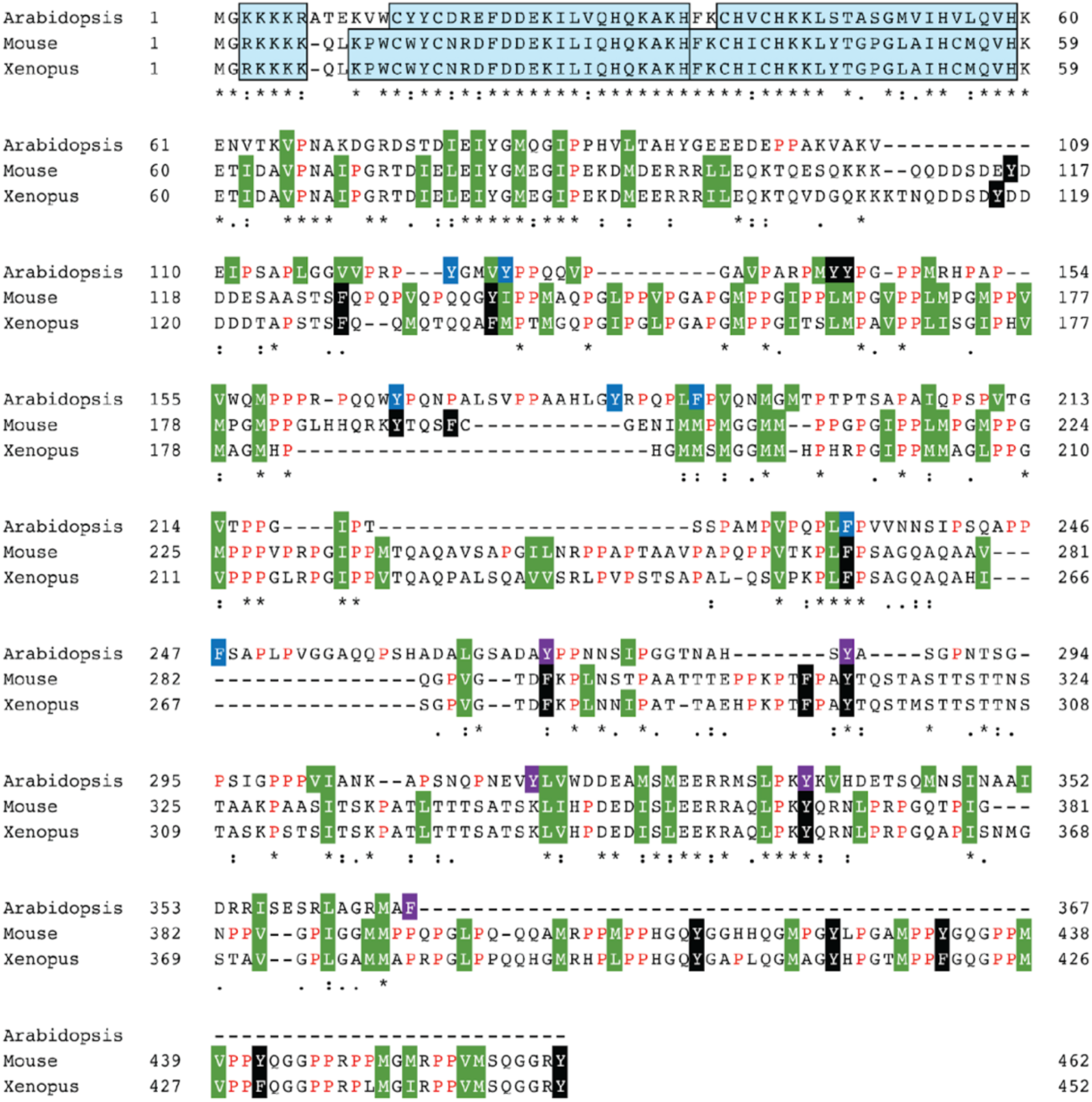
Sequence alignment of *Arabidopsis* SUF4, mouse and Xenopus BuGZs. N-terminal domain that contains predicted NLS and zinc fingers (highlighted in cyan), is conserved well among three SUF4/BuGZ sequences whereas the central/C-terminal disordered domain of *Arabidopsis* SUF4 seems diverged from animal BuGZs. F and Y residues in the predicted disordered region are highlighted in black, blue, or purple. Other strong hydrophobic residues are highlighted in green. Among F and Y in the predicted disordered region of SUF4, the last 5 F and Y (highlighted in purple) were mutated to S in the 5S mutant and additional 7 F and Y at the central region (highlighted in blue) were mutated to the S in the 12S mutant.

**Supplemental Figure 4.**
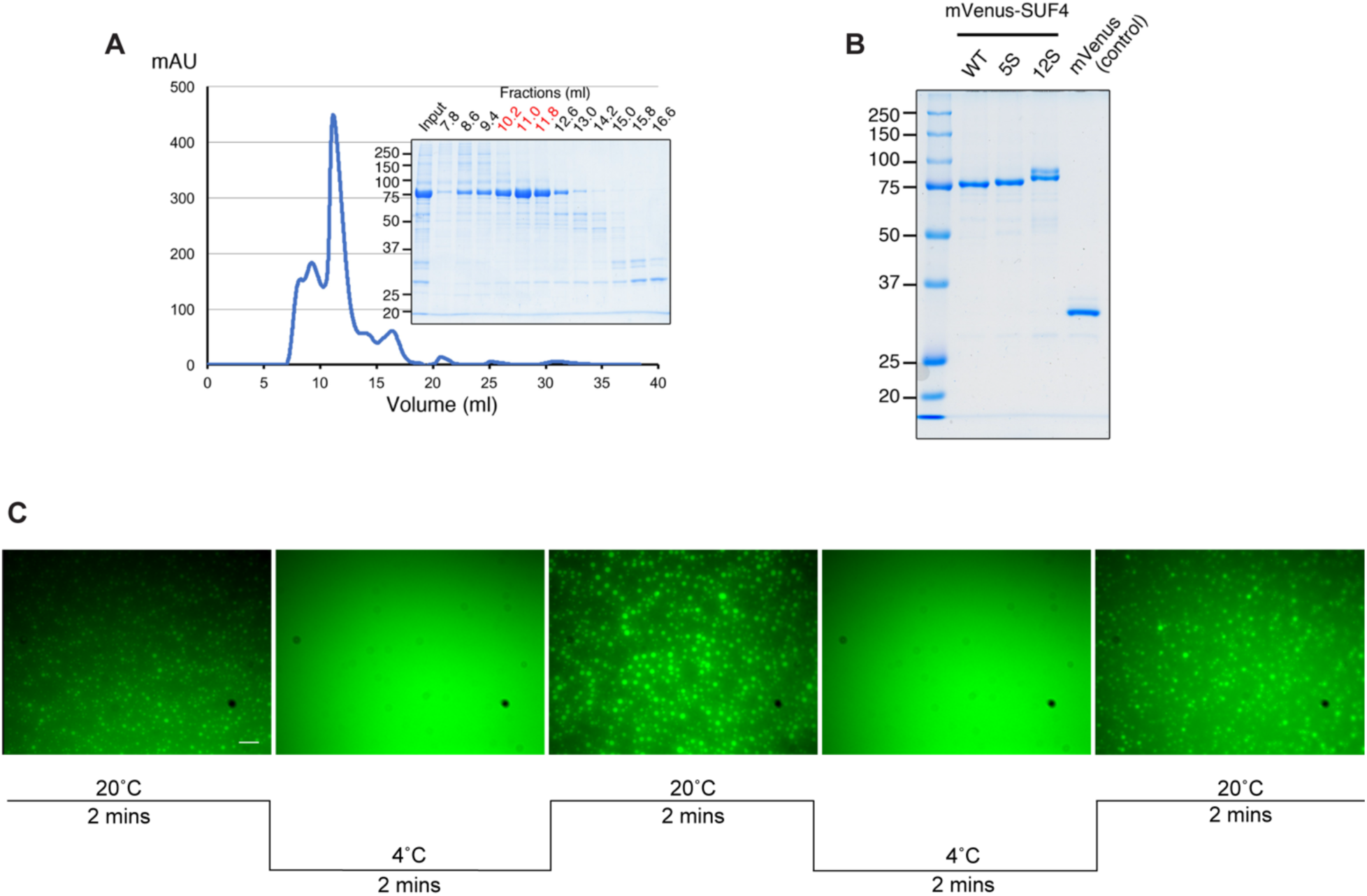
Purified SUF4 forms round and reversible liquid-like condensates *in vitro*. **A)** A representative elution profile of His-mVenus-SUF4 on gel filtration chromatography. Recombinant proteins were expressed in Sf9 insect cells, purified with IMAC, followed by gel filtration. Fractions were eluted around 11-ml after sample loading to enrich for the fusion proteins (indicated with red letters in the inset of the gel image of fractions). **B)** SDS-PAGE analysis of the purified His-mVenus-SUF4 and control HIS-mVenus proteins. Note that 12S mutant protein has two bands on gel. **C)** Purified SUF4 forms round and reversible liquid-like condensates *in vitro*. SUF4 can undergo multiple rounds of temperature shifts without transitioning into a more stable gel or solid-like aggregate. Prior to temperature shifts, SUF4 is incubated at either 20°C to 4°C for 2 minutes. Upon temperature shifts every two minutes, SUF4 changes its phase state and maintains this behavior until the next shift. Scalebar: 45µm.

**Supplemental Figure 5.**
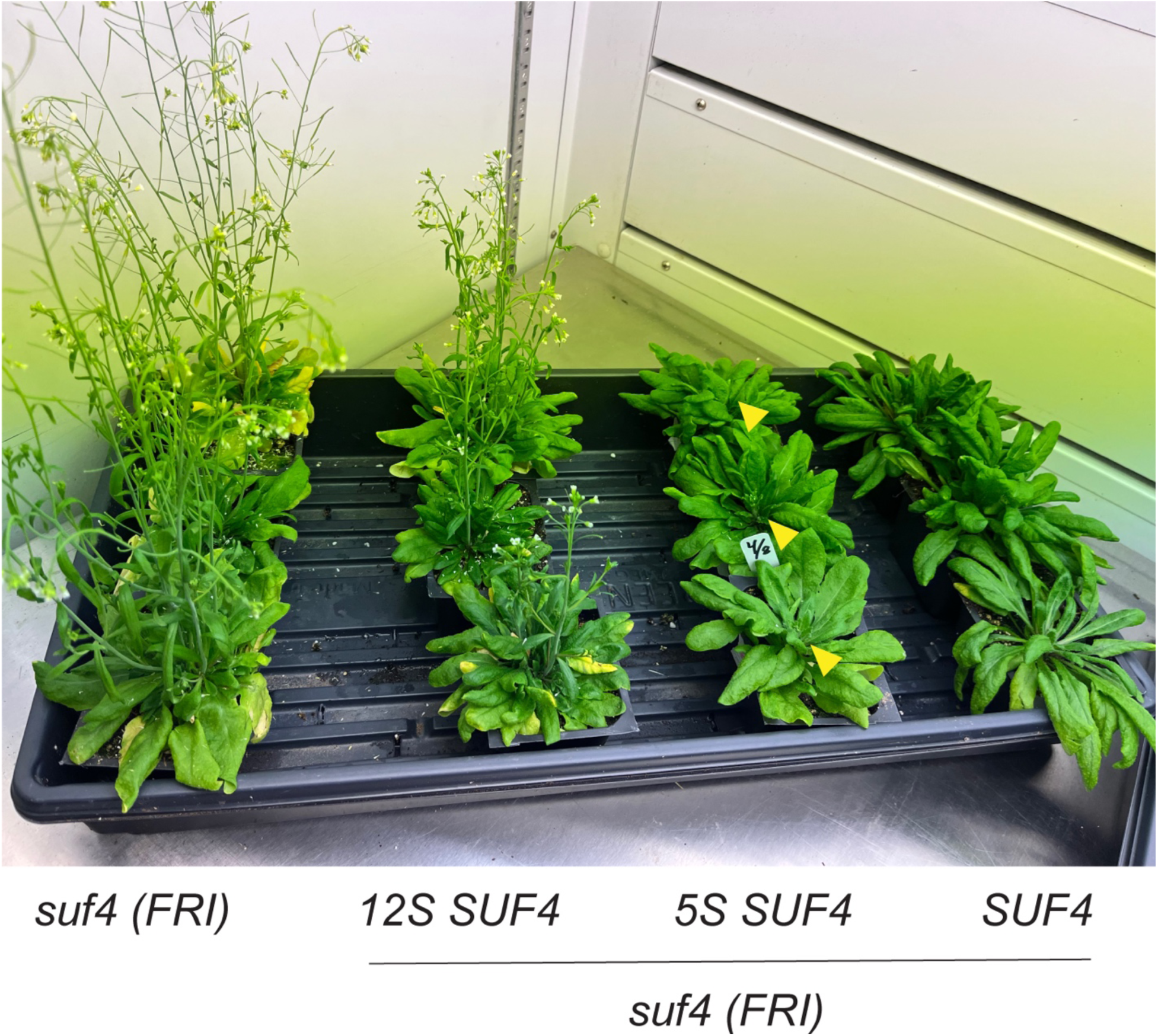
Image of SUF4 variants in comparison to mutant *suf4*, which flowers early. Left to right: *suf4 (FRI)* mutant with an early flowering time phenotype, 12S SUF4 (12 F and Y substitutions for S) with a moderate early flowering phenotype, 5S SUF4 (5 Y and F substitutions for S) with weak early flowering phenotype, rescue mEGFP-SUF4 (functional copy of SUF4) rescuing the delayed flowering time phenotype.

**Supplemental Figure 6.**
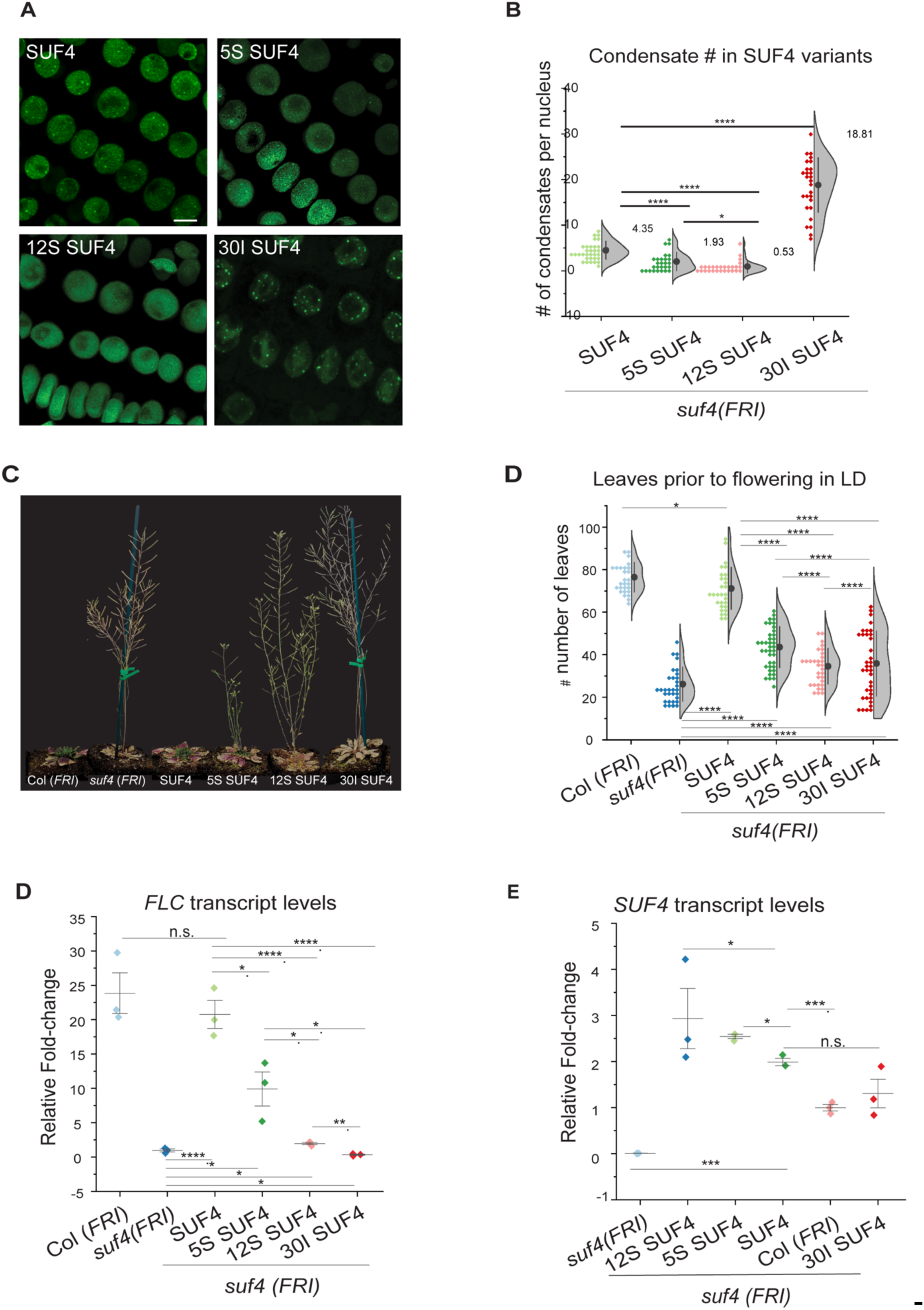
Characterization of 30I SUF4 in relation to the remaining SUF4 variants. **A)** Confocal images showing that 30I SUF4 assembles into more nuclear condensates than SUF4 rescue. Scale bar: 5µm. **B)** quantification of A). 30I SUF4 forms an average of ∼18 condensates per nucleus). **C-D)** Leaf counting assay of different variants in long- and short-day conditions. Note that 30I SUF4 does not rescue the flowering time phenotype. Statistical analysis done using a Mann-Whitney U test. **E)** Relative abundance of *FLC* transcripts in 30I compared to different SUF4 variants. 30I SUF4 does not rescue *FLC* transcript levels. **F)** qPCR of *SUF4* in 30I and other transgenic *SUF4* variant lines. 30I SUF4 exhibits similar levels of transcripts as the rescue SUF4 line. Statistical analysis for E) and F) conducted using a two-tailed Student’s t-test for independent means. Error bars represent the standard deviation. All data other than 30I SUF4 are reproduced from the main figures. Asterisks denote P-values * <0.01, ** <0.001, ***<0.001, and ****<0.0001

**Supplemental Figure 7.**
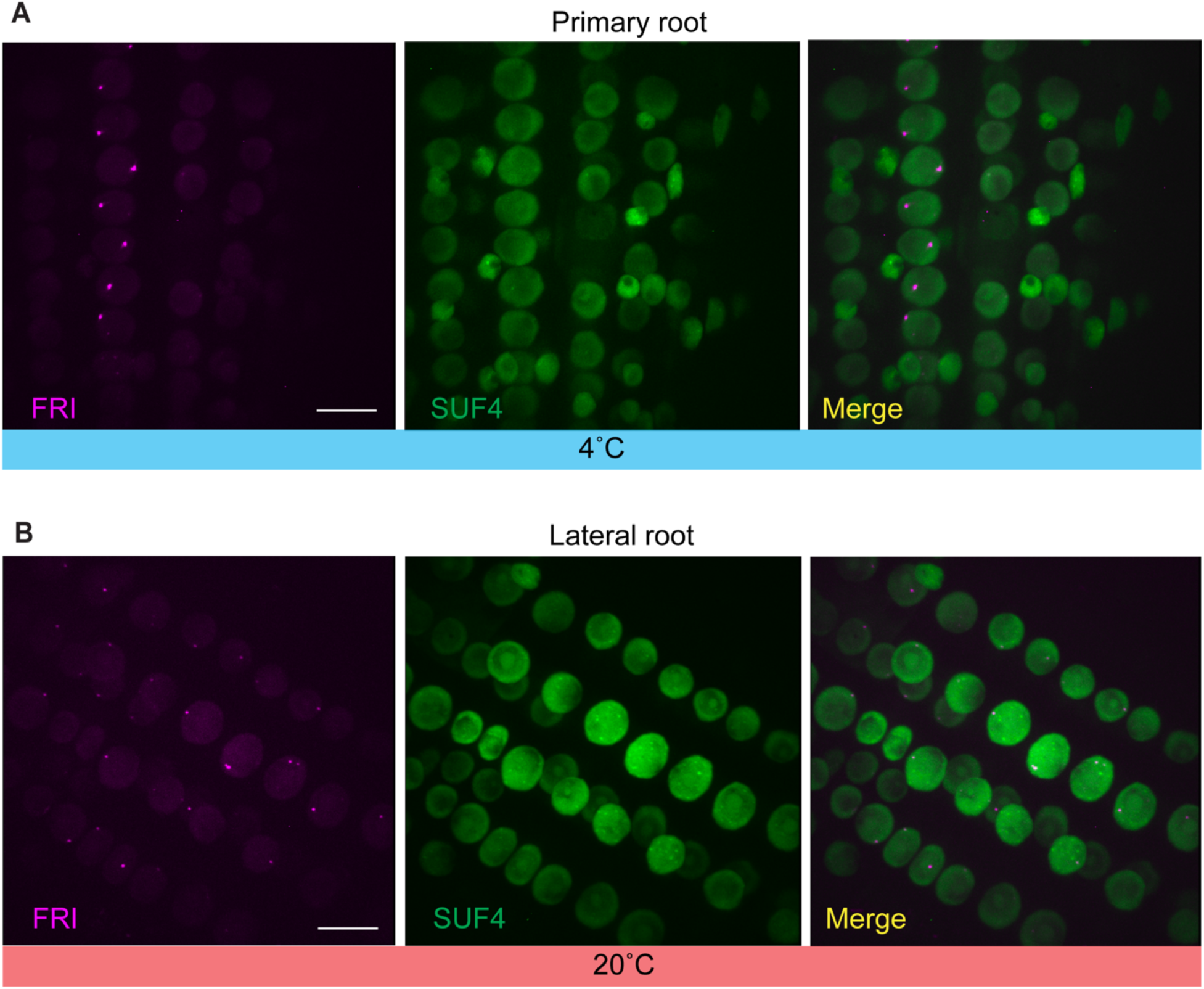
**A)** FRI forms condensates in the primary root of 6 day old seedlings after 24 hours of 4°C treatment. **B)** FRI forms condensates in a burst-like fashion in lateral roots under warm temperatures. Scalebars:10µm

## Acknowledgements

We would like to thank Carlos Castañeda, Broder Schmidt, Adrienne Roeder, Rick Amasino, Doris Wagner, Ryan O’Hanlon, as well as the members of the Ehrhardt lab, IDPSIG, Carnegie Institution for Science, and Stanford University for their insightful comments and meaningful discussions.

## Funding

H.M.M., T.H., Y.Z., and D.W.E. was supported on the Carnegie Venture Grant Fund. H.M.M. was also supported by the Simons Foundation-Life Science Research Foundation [Meyer LSRF18/Carnegie Fund 10861].

## Author contributions

Conceptualization: HMM, DWE, YZ

Methodology: HMM, DWE, YZ, TH, AM

Investigation: HMM, TH, DWE

Visualization: HMM, DWE

Funding acquisition: HMM, DWE, YZ

Supervision: DWE, YZ

Writing – original draft: HMM, DWE

Writing – review & editing: HMM, DWE, YZ

## Competing interests

Authors declare that they have no competing interests

## Data and materials availability

All data are available in the main text or the supplementary materials

## Notes

### Competing Interest Statement

The authors have declared no competing interest.

### Summary of Updates

The text and figures have been revised for clarity and for journal submission

